# Systematic silencing of oncogenic fusion transcripts with ultra-precise CRISPR-Cas13b

**DOI:** 10.1101/2025.02.10.637348

**Authors:** Wenxin Hu, Honglin Chen, Joshua ML Casan, Carolyn Shembrey, Steven He, Lauren M Brown, Timothy P. Hughes, Deborah L. White, Antoine De Weck, Ilia Voskoboinik, Joseph A Trapani, Paul G Ekert, Teresa Sadras, Mohamed Fareh

**Affiliations:** Peter MacCallum Cancer Centre, Melbourne, 3000, Australia; Sir Peter MacCallum Department of Oncology, The University of Melbourne, Parkville, 3052, Australia; Children’s Cancer Institute, Lowy Cancer Research Centre, UNSW Sydney, NSW Australia 2052; School of Women’s and Children’s Health, UNSW Sydney, Sydney, NSW, Australia; Precision Cancer Medicine Theme. South Australian Health and Medical Research Institute (SAHMRI). Adelaide SA; Faculty of Health and Medical Sciences, University of Adelaide, Adelaide, SA 5005, Australia; Murdoch Children’s Research Institute, Royal Children’s Hospital, 50 Flemington Rd, Parkville, Melbourne, 3052, Australia

**Keywords:** CRISPR-*Psp*Cas13b, Oncogenic fusion transcripts targeting, Drug-resistance, Personalised cancer therapy

## Abstract

Oncogenic gene fusions are key drivers of cancer, yet most remain untargetable by current therapies. Here, we establish CRISPR-*Psp*Cas13b as a personalizable platform for systematic silencing of various fusion transcripts. We reveal that recognition and cleavage of the breakpoint sequence by PspCas13b disrupts the fusion transcript, resulting in unexpected RNA nicking and ligation near the cleavage site, which generates out-of-frame, translation-incompetent transcripts. This approach efficiently degrades canonical and drug-resistant BCR::ABL1 mutants (e.g., T315I), a primary cause of resistance to tyrosine kinase inhibitors (TKIs) and relapse in chronic myeloid leukemia (CML). Silencing T315I BCR::ABL1 mRNA in drug-resistant CML cells triggers extensive transcriptomic, proteomic, and phosphoproteomic remodelling, causing erythroid differentiation and apoptosis. Beyond BCR-ABL1 mutants, personalized design of *Psp*Cas13b effectively silences other undruggable fusions, including RUNX1::RUNX1T1 and EWSR1::FLI1, key drivers in acute myeloid leukemia and in Ewing sarcoma, respectively. Collectively, this study establishes a framework for systematic, precise, and personalizable targeting of otherwise *undruggable* or drug-resistant oncogenic transcripts.

## INTRODUCTION

Recent advances in next-generation sequencing have enabled rapid identification of oncogenic transcripts in individual patients within actionable time frames^1–3^. Numerous fusion genes generated by chromosomal translocations demonstrate potent oncogenic activity but also remain largely ‘*undruggable*’ with conventional therapeutics^4^. Identified fusion genes contribute to approximately 20% of all human cancer cases, though their prevalence varies significantly across cancer types. In some cancers, the frequency is particularly high, with around 62% of leukemias and 85% of Ewing’s sarcomas exhibiting gene fusion alterations^6^. Fusion-driven cancers typically exhibit a low tumour mutational burden, with the fusion gene as the primary driver and minimal additional mutations. Targeting these fusions may produce a significant therapeutic effect, as they are central to tumorigenesis^7^. However, personalised targeting of these fusion variants at the protein level has proven to be challenging due to structural complexity^8–10^, and in the rare cases where small molecule inhibitors are available, the rapid development of drug resistance and consequent disease relapse is common^11,12^. Additionally, recent studies indicate that blocking the catalytic activity (kinase domain) of certain gene fusions with small molecules is often insufficient to fully suppress their oncogenic activity^13^. In fact, the scaffold functions of these gene fusions can also drive oncogenesis, suggesting that targeted protein or RNA degradation strategies may provide superior therapeutic efficacy^13^.

Notably, the unique chimeric mRNA sequence encoded at the fusion site of the two genes represents a tractable and potentially impactful target for sequence-specific silencing with programmable RNA nucleases.

The type VI CRISPR (clustered regularly interspaced short palindromic repeats) effectors termed CRISPR-Cas13 (Cas13) are programmable RNA-guided targeting enzymes that exclusively degrade single-stranded RNA with high efficacy and specificity^14,15^. Recent studies have deployed Cas13 systems in a variety of targeted RNA manipulations, including RNA silencing^5,16^, nucleic-acid detection^17–19^, precise RNA base editing^20,21^, and viral suppression^22–24^. The efficiency and transience of RNA targeting with Cas13 represents a promising approach to specifically edit oncogenes without risking permanent alteration of the genome in somatic and germline cells; an inherent limitation of DNA-editing CRISPR enzymes^25–27^. Therefore, Cas13 is highly attractive for targeting aberrant fusion transcripts that drive various human genetic diseases including cancer. Among various characterised Cas13 enzymes, *Psp*Cas13b ortholog possesses high silencing efficiency and specificity, underscoring its suitability for targeted gene silencing in human cells^5,28^.

Here, we show that the personalised design of *Psp*Cas13b targeting the breakpoint sequence enables specific and potent silencing of various ABL1-related fusion transcripts, including *BCR::ABL1*, a master driver of chronic myeloid leukemia (CML) and acute lymphoblastic leukemia (ALL)^29,30^. Targeting the breakpoint suppressed both the canonical BCR::ABL1 and TKI-resistant mutants (T315I), which frequently drives treatment failure and disease progression in CML patients^31^. The effective silencing of T315I BCR::ABL1 mutant in drug-resistant CML cells led to profound transcriptomic and proteomic remodelling, erythroid differentiation and apoptosis. Beyond ABL1-fusions, we reprogramed *Psp*Cas13b to precisely recognise and cleave other undruggable fusion transcripts. Thus, this study provides a framework for efficient and specific suppression of various undruggable or drug-resistant fusion transcripts through personalised targeting of the breakpoint sequence.

## RESULTS

### Efficient Silencing of Oncogenic Fusion Drivers with Personalized Design of ***Psp*Cas13b.**

We investigated whether *Psp*Cas13b can be reprogrammed to silence oncogenic fusion gene transcripts. We hypothesised that the breakpoint at the interface of the two partner genes offers a unique targetable sequence at the RNA level that may allow selective silencing of fusion transcripts. To test this, we designed crRNAs targeting the breakpoint of four oncogenic ABL1 fusion genes, BCR::ABL1-p190 (BCR exon1 to ABL1 exon2 translocation), BCR::ABL1-p210 (BCR exon14 to ABL1 exon2 translocation), SFPQ::ABL1, and SNX2::ABL1, which are all established drivers of ALL or CML. These fusion genes involve the rearrangement of *ABL1* with a partner gene, leading to a constitutively active ABL1 tyrosine kinase domain^29,32–34^. The fusion genes were each cloned into an IRES-eGFP reporter construct that produces a gene-of-interest-IRES-eGFP chimeric transcript, which is subsequently translated into separate proteins due to the presence of the IRES sequence. If the chimeric RNA is cleaved by *Psp*Cas13b at the targeted site, the resultant RNA fragments lose key features such as their 5’cap, 5’/3’UTRs, and polyA tail, impairing their intracellular stability and/or their translation. As a result, both the cleaved fusion gene sequence and the downstream reporter gene (eGFP) undergo degradation and/or translational repression (**Supplementary Fig. 1**). Therefore, efficient cleavage of the fusion transcript by *Psp*Cas13b can be inferred from the loss of eGFP fluorescence.

**Supplementary Figure 1.**
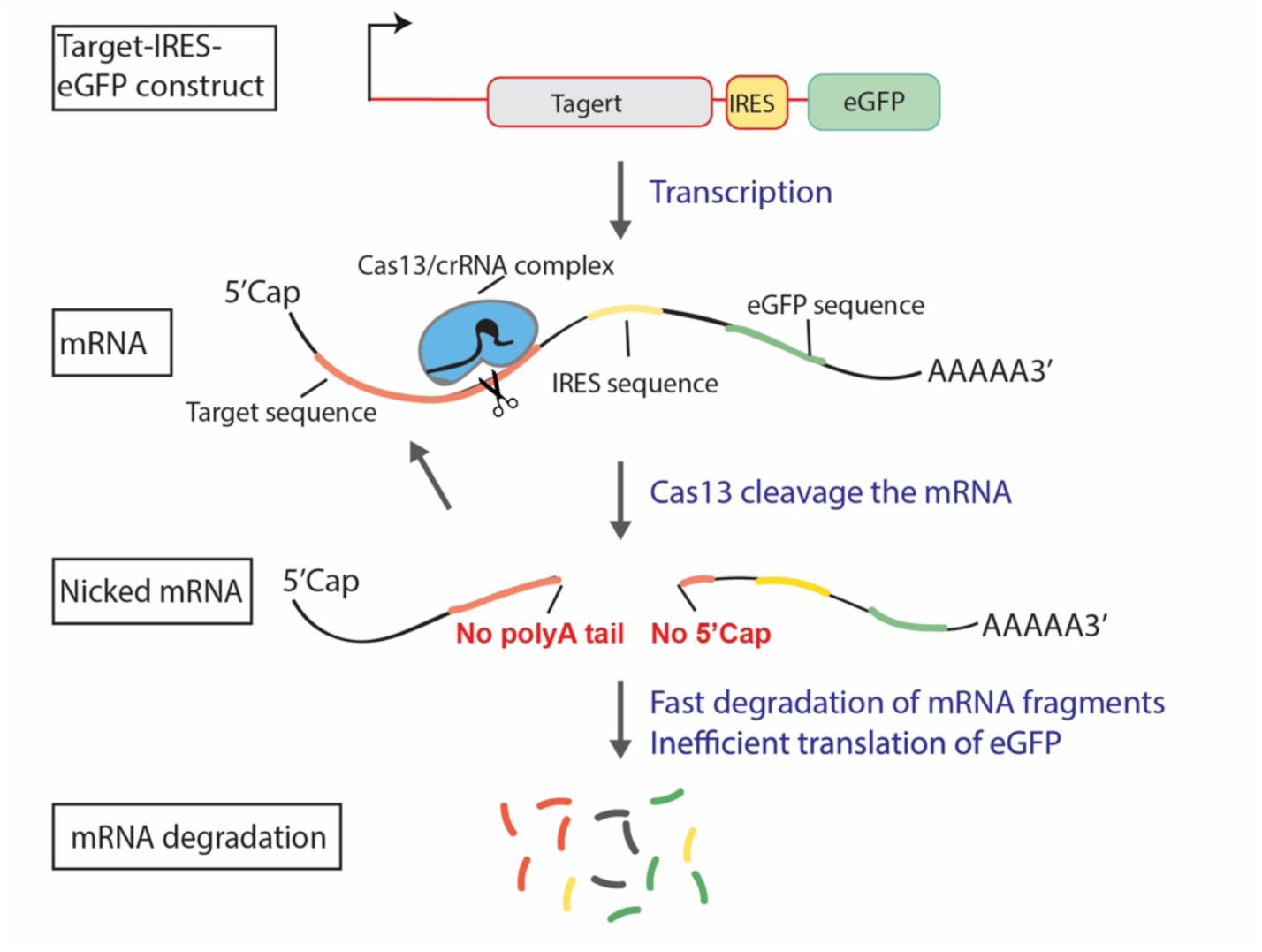
Schematic of fluorescent reporter assay used to track degradation of various fusion transcripts. The fusion genes were each cloned into an IRES-eGFP reporter construct, which is subsequently transcribed into a single chimeric RNA encoding a fusion protein, IRES sequence, and eGFP. *Psp*Cas13b cleaves the chimeric RNA at the fusion gene breakpoint. The RNA fragments lose key RNA features like their 5’cap, 5’/3’UTRs, and polyA tail, and therefore undergo RNA degradation and/or inefficient protein translation.

We designed 9 unique crRNAs tiled across the breakpoint of each ABL1 fusion gene, in which the binding sites of consecutive crRNAs are spaced by 3 nucleotides. Each crRNA was transfected into HEK 293T cells together with *Psp*Cas13b and the ABL1 fusion-gene-IRES-eGFP plasmids. We observed the ∼80 to 90% silencing of all four fusion genes via fluorescence microscopy and confirmed these results by using RT-qPCR, although the efficiency varied depending on crRNA binding sites (**Figure 1a-1g**). We further confirmed high silencing of BCR::ABL1-p190 at the protein level by western blot analysis, with -12, -9 and +12 crRNAs exhibiting the highest silencing efficiencies (**Figure 1h)**. Analysis of STAT5 and ERK phosphorylation, hallmarks of BCR::ABL1 dependent kinase activity (**Figure 1i**), revealed that potent crRNAs can efficiently suppress BCR::ABL1 and its downstream oncogenic networks (**Figure 1j**). Similarly, 1 μM imatinib, a small inhibitory molecule that blocks the tyrosine kinase domain of ABL1 (**Figure 1i**), inhibited BCR::ABL1 mediated phosphorylation of STAT5 and ERK without altering the expression levels of BCR::ABL1 protein, whereas *Psp*Cas13b crRNAs efficiently silenced BCR::ABL1 protein expression (**Figure 1j**). Interestingly, the most potent crRNA+12 showed greater suppression of STAT5 phosphorylation, consistent with its high efficacy in depleting the BCR::ABL1 protein through mRNA silencing (**Figure 1j**). In conclusion, these findings demonstrate that personalised design of *Psp*Cas13b to efficiently silence fusion transcripts through the recognition of their unique breakpoint sequence.

**Figure 1.**
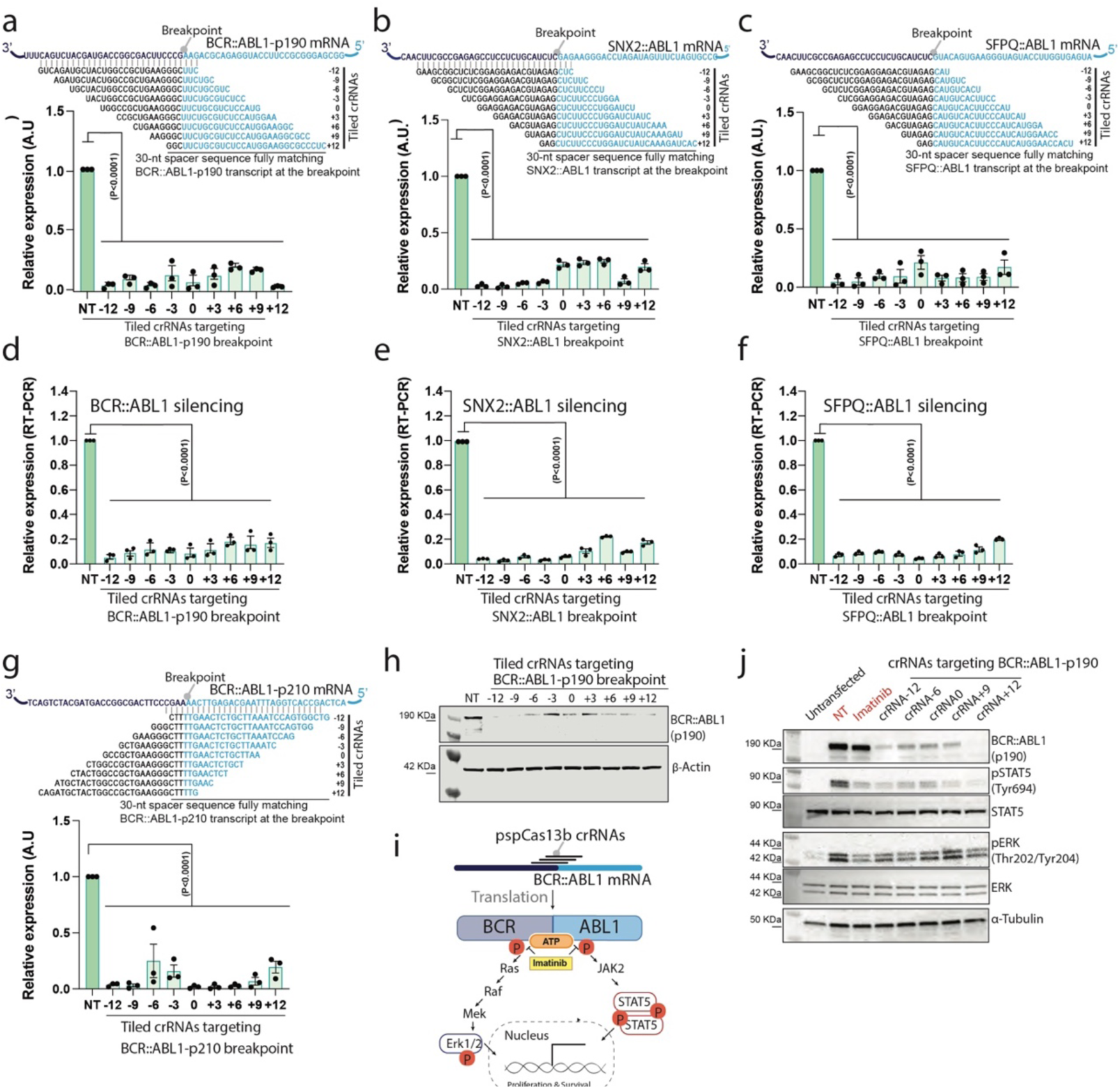
Reprogrammed *Psp*Cas13b suppresses fusion gene transcripts with high efficiency. **(a-c, g)** EGFP reporter assay to assess the silencing efficiency of tiled *Psp*Cas13b crRNAs with 3-nucleotide resolution targeting the breakpoint region of fusion transcripts BCR::ABL1-p190 **(a)**, SNX2::ABL1 **(b)** SFPQ::ABL1 **(c) and (g)** BCR::ABL1-p210. Data points in the graphs are averages of mean fluorescence from 4 representative fields of view per condition imaged; *N* = 3. The data are represented in arbitrary units (A.U.). Errors are SEM and *p*-values of the one-way ANOVA test are indicated (95% confidence interval). **(d-f)** RT-PCR assays measuring silencing efficiency of tiled *Psp*Cas13b crRNAs (3-nucleotide resolution) targeting the breakpoint regions of fusion transcripts **(d)**, BCR::ABL1-p190, **(e)** SNX2::ABL1 and **(f)** SFPQ::ABL1; *N* = 3; Data are normalised means and errors are SEM; Results are analysed by one-way ANOVA test with *p*-values indicated (95% confidence interval). **(h)** Representative Western blot analysis to examine the expression level of BCR::ABL1-p190 protein in HEK 293T cells expressing tiled crRNAs targeting BCR::ABL1-p190 transcripts 24 h post-transfection; *N* = 3. (See uncropped blots in the Source file). **(i)** Schematic of BCR::ABL1 dependent phosphorylation of ERK and Stat proteins, and inhibition of BCR::ABL1 oncogenic activity with imatinib, a potent competitor of ATP binding to ABL kinase. **(j)** Representative Western blot analysis examining the suppression of BCR::ABL1 expression and the subsequent inhibition of STAT5 and ERK phosphorylation in HEK 293T cells expressing BCR::ABL1-p190, *Psp*Cas13b and either NT or crRNA targeting the BCR::ABL1 24 h post-transfection. HEK 293T cells expressing BCR::ABL1-p190 and *Psp*Cas13b treated with 1µM imatinib for 4 hours were used as a positive control. Parental cells are HEK 293T cells transfected with *Psp*Cas13b, NT and a random control plasmid. This condition shows the baseline expression of pSTAT5 and pERK in a BCR::ABL1 independent manner; *N* = 2. *N* is the number of independent biological experiments. Source data are provided as a Source data file.

### *Psp*Cas13b Discriminates Between Fusion and Wildtype Variants Despite Their Extensive Sequence Homology

Our previous spacer mutagenesis study indicated that *Psp*Cas13b can discriminate between RNAs that share extensive (∼80%) sequence homology^5^. We investigated whether this targeting resolution can be generalised to oncogenic fusion RNAs. To determine this, we introduced increasing non-consecutive nucleotide mismatches between the spacer of BCR::ABL1-p190 crRNA (crBCR::ABL1) and the target breakpoint sequence (**Figure 2a**). The data revealed that up to three nucleotide mismatches were well tolerated. However, four or higher numbers of non-consecutive nucleotide mismatches drastically impaired crRNA silencing efficiency (**Figure 2a**). Having six consecutive nucleotide mismatches led to a notable loss of silencing when situated at the 3’ end (spacer positions 25-30) or the central region (spacer positions 12-17), while 9 consecutive nucleotide mismatches dramatically reduced silencing irrespective of position (**Figure 2b**). Western blot analysis of BCR::ABL1 protein expression confirmed these data and showed that 3-nucleotide mismatches are well tolerated, while 4-nucleotide mismatches or higher led to substantial or complete loss of silencing (**Figure 2c**). This data indicates that mismatch tolerance is strongly influenced by the number of unpaired bases and their positions within the spacer. To maximize specificity, the breakpoint should be positioned near the center of the spacer to minimize off-target effects on untranslocated fusion partners, such as wild-type BCR or ABL1. To confirm this specificity, we tested crRNAs targeting BCR::ABL1 fusion against wildtype untranslocated BCR and ABL1 transcripts which are usually expressed in normal tissues. We cloned constructs encoding partial mRNA sequences of the BCR::ABL1 fusion, ABL1 alone, and BCR alone in frame with mCherry, eGFP, or TagBFP fluorescent reporters, respectively (**Figure 2d-2f**). We designed 3 crRNAs targeting the BCR::ABL1 breakpoint sequence (crBCR::ABL1), BCR sequence (crBCR), or ABL1 sequence (crABL1) that we tested against the aforementioned constructs. The fluorescence signals from mCherry, eGFP, and TagBFP enable accurate quantification of on-target and off-target silencing with these crRNAs. As anticipated, all 3 crRNAs silenced the *bona fide* BCR::ABL1 transcript as this mRNA possesses full-length spacer binding sites for all three crRNAs (**Figure 2d, Supplementary Fig. 2a**). However, ABL1 and BCR transcripts were silenced only by their cognate crABL1 and crBCR crRNAs (**Figure 2e & 2f, Supplementary Fig. 2a**). Notably, crBCR::ABL1 targeting the breakpoint sequence did not affect either BCR or ABL1 wildtype transcripts due to the 15-nucleotide mismatch (**Figure 2e & 2f**). Western blot analysis confirmed these findings (**Figure 2d-2f**), demonstrating *Psp*Cas13b specifically silences oncogenic fusion transcripts without off-targeting untranslocated transcripts.

**Figure 2.**
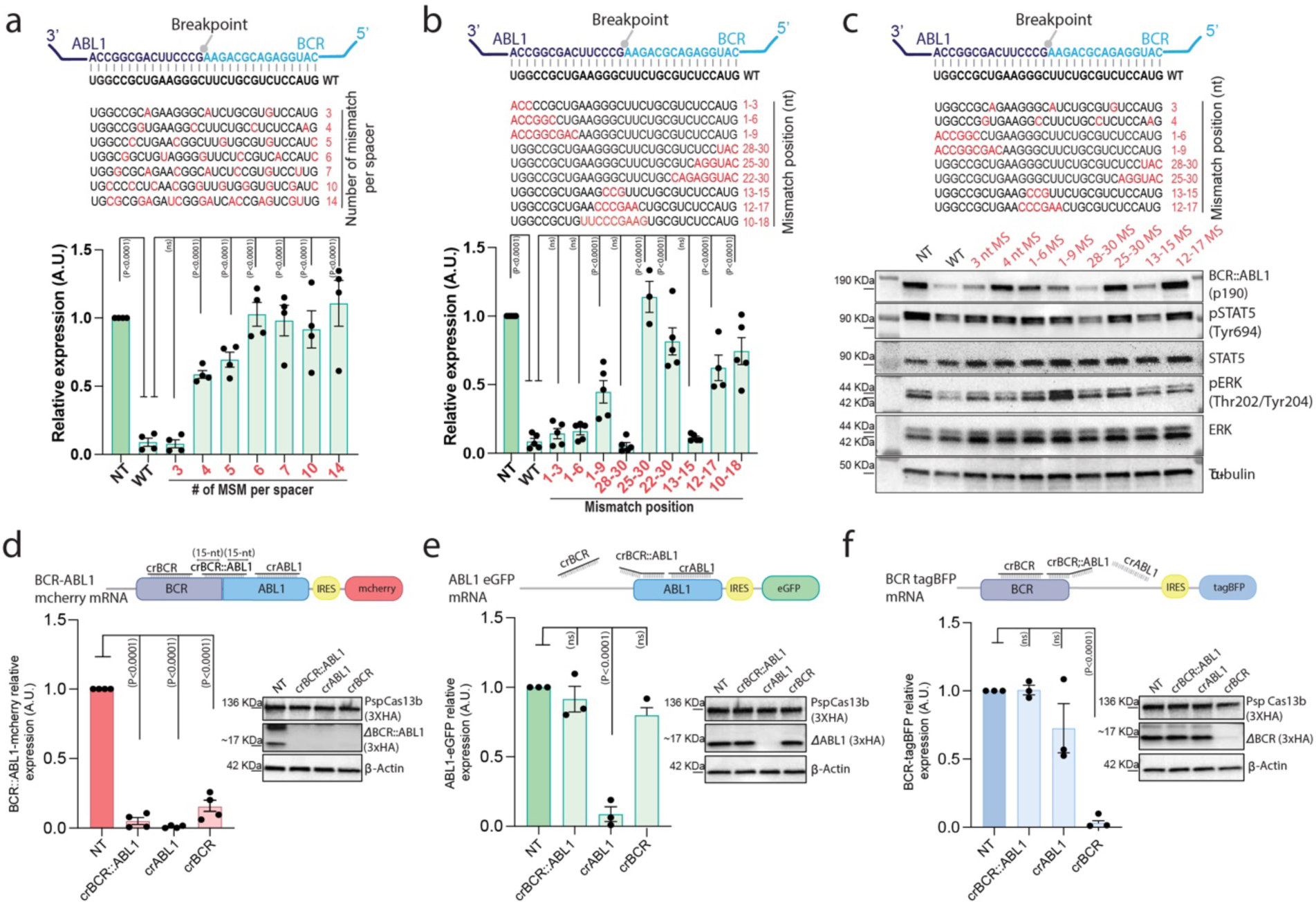
The targeting of the breakpoint can efficiently discriminate between translocated tumour RNAs and wildtype variants despite extensive sequence homology. **(a–b)** Comprehensive analysis of spacer-target interaction examining specificity and mismatch tolerance of *Psp*Cas13b crRNAs targeting the breakpoint region of BCR::ABL1-p190 transcript. The nucleotides in red highlight various mismatch positions in the spacer sequence. Data points in the graph are averages of mean fluorescence from 4 representative fields of views per condition imaged; *N* = 4. The data are represented in arbitrary units (A.U.). Errors are SEM and *p*-values of the one-way ANOVA test are indicated (95% confidence interval). **(c)** Western blot analysis examining the expression level of BCR::ABL1-p190 protein and phosphorylation status of STAT5 and ERK in HEK 293T cells expressing crRNAs with various mismatches 24 h post-transfection; *N* = 3. (See uncropped blots in the Source file). **(d-f)** Three colour fluorescence-based reporter assays to assess the specificity of crRNA targeting the breakpoint region of BCR::ABL1 in HEK 293T cells 48 h post-transfection. The schematics show **(d)** BCR::ABL1-mCherry mRNA, **(e)** ABL1-eGFP mRNA, and **(f)** BCR-TagBFP mRNA and their interaction with crBCR, crBCR::ABL1 and crABL1 crRNAs through full, partial, or no spacer-target basepairing. The quantifications of the silencing of **(d)** BCR::ABL1-mCherry, **(e)** ABL1-eGFP and **(f)** BCR-TagBFP. Data points are averages of normalised mean fluorescence from 4 representative fields of view per condition imaged. The data are represented in arbitrary units (A.U.). Errors are SEM and *p*-values of the one-way ANOVA test are indicated (95% confidence interval). *N* = 3. The right panels show representative western blot analyses to examine the expression levels of ΔBCR-ABL, ΔABL1, and ΔBCR proteins from data in d-f; *N* = 3 (See uncropped blots in the Source file).

### Transcriptomic Analysis Confirms the High Specificity of *Psp*Cas13b and Reveals an Unexpected Pattern of Target Cleavage

Several Cas13 orthologs have been shown to possess collateral cleavage activity, in which target recognition can trigger the indiscriminate degradation of nearby cellular RNAs. While this collateral activity is well studied in biochemical *in-vitro* assays^18^ and bacteria^15^, the extent to which it occurs in mammalian cells remains controversial^35,36^. We found that upon BCR::ABL1 target recognition and nuclease activation in HEK 293T, *Psp*Cas13b did not cleave 28S RNA. In contrast, the *Rfx*Cas13d ortholog exhibited collateral activity in that it partially nicked the ribosomal 28S RNA (**Supplementary Fig. 2b**), which is consistent with findings in a previous study^28^. This suggested that, unlike *Rfx*Cas13d, *Psp*Cas13b possesses no or very limited collateral activity in mammalian cells.

To better determine the impact of *Psp*Cas13b nuclease activation on global endogenous cellular RNA, we performed RNA sequencing on HEK 293T cells expressing BCR::ABL1 and *Psp*Cas13b. The transcriptomic data shows no significant downregulation of endogenous transcripts due to off-target or collateral effects in HEK 293T cells (**Figure 3a**). However, standard short read RNA sequencing analysis showed no significant reduction in BCR::ABL1 mRNA levels (**Figure 3b**, **Supplementary Fig. 2c).** This was surprising as it contradicted western blot and RT-qPCR data shown above (**Figure 1d-1h**). When we carefully examined the fate of the RNA regions flanking the *Psp*Cas13b binding site, we noticed that a ∼500nt long sequence surrounding the breakpoint was largely trimmed, while distal regions to the cleavage site remained stable (**Figure 3c**).

**Figure 3.**
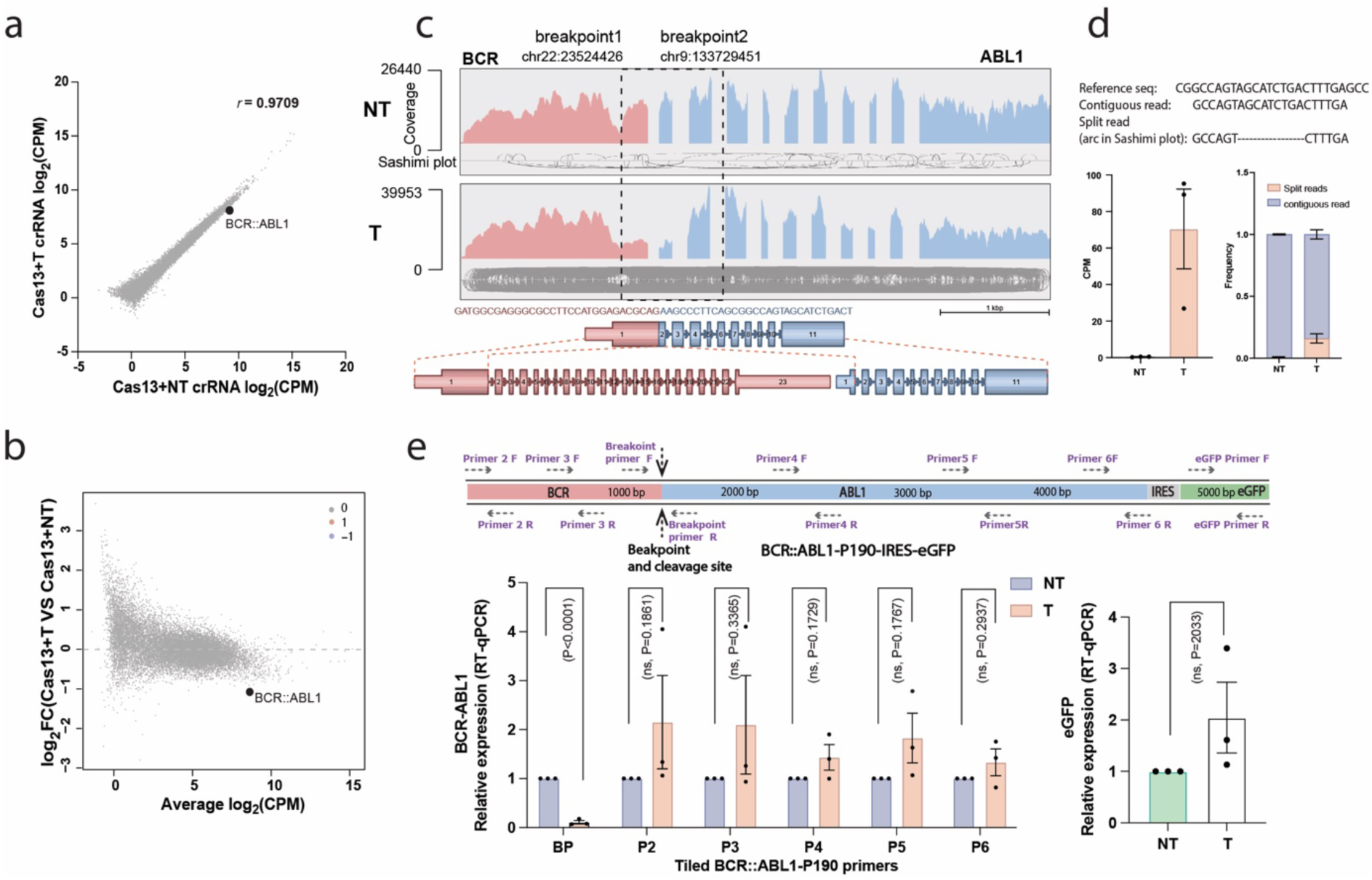
Transcriptomic analysis confirms the high specificity of *Psp*Cas13b and reveals an unexpected pattern of target cleavage. RNA sequencing assays were performed in HEK 293T cells transfected with plasmids coding *Psp*Cas13b, BCR::ABL1-p190-IRES-eGFP and either a crRNA targeting the breakpoint of BCR::ABL1-p190 transcript (T) or non-targeting (NT) crRNA. RNA was extracted 48 hours post-transfection. **(a)** A linear regression plot showing log_2_-counts per million (CPM) of transcripts and **(b)** a mean-difference plot (MD-plot) showing the log_2_-fold change(FC) and average abundance of transcripts in HEK 293T cells expressing *Psp*Cas13b with a T crRNA versus an NT crRNA. Each data point represents a gene, and genes with log_2_FC > 1 (upregulation, indicated in red) or log_2_FC < -1 (downregulation, indicated in blue), and p-value < 0.05 are considered as significant differentially expressed transcripts. *N*=3; (*N* represents the number of biological replicates). **(c)** Representative Arriba fusion plots showing the read alignments against the BCR::ABL1 transcript in HEK 293T cells expressing *Psp*Cas13b with either an NT (top) or a T (bosom) crRNA. The breakpoint region is highlighted with a dashline box. **(d)** Counts per million (CPM) (lel) and frequency (right) of split reads exclusively mapped to BCR::ABL1 breakpoint (filter criteria shown in the method section) in HEK 293T cells treated with *Psp*Cas13b or imatinib. **(e)** Schematic showing the design of RT-qPCR primers binding different regions on BCR::ABL1-p190-IRES-eGFP transcript to determine the fate of cleavage RNA fragments (top). RT-qPCR assays (bosom) to measure the expression levels of BCR::ABL1-p190 and eGFP using primers shown in the top schematic. *N*=3. Data are normalised means and errors are SEM. Results are analysed by unpaired two-tailed Student’s t-test (95% confidence interval). *N* represents the number of biological replicates). Source data are provided as a Source data file.

**Supplementary Figure 2.**
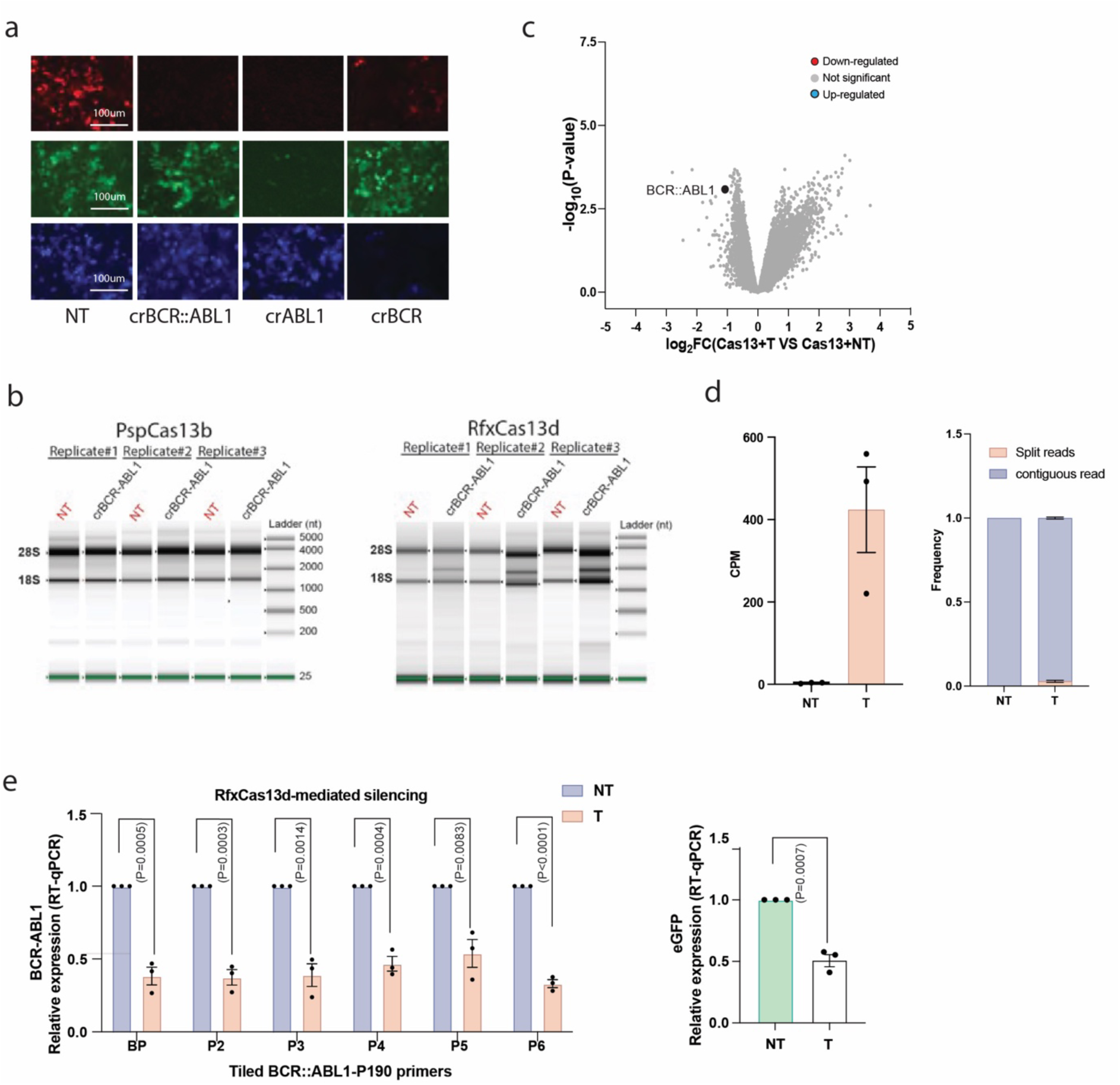
**(a)** Representative fluorescence microscopy images show the silencing efficiency of crBCR, crBCR::ABL1 and crABL1 crRNAs targeting BCR::ABL1-mCherry, ABL1-eGFP and BCR-TagBFP transcripts in HEK 293T cells. Scale bar = 100 μm, *N* = 3. Unprocessed representative images are provided in the Source Data file. **(b)** Assessment of potential *Psp*Cas13b (lel) and *Rfx*Cas13d (right) collateral activity through the analysis of ribosomal RNA (28S and 18S) degradation in the presence of target-free (NT) or target-bound (crBCR::ABL1) Cas13 nuclease, *N*=3. *N* is the number of independent biological experiments. **(c)** A volcano plot of transcripts showing the log_2_-fold change (FC) in HEK 293T cells expressing *Psp*Cas13b with a T crRNA versus an NT crRNA. Each data point represents a gene, and genes with log_2_FC > 1 (upregulation, indicated in red) or log_2_FC < -1 (downregulation, indicated in blue), and p-value < 0.05 are considered as significant differentially expressed transcript, *N*=3. **(d)** Counts per million (CPM) of split reads that mapped to BCR::ABL1 transcript in HEK 293T cells treated with *Psp*Cas13b or imatinib (lel). The frequency of split and contiguous reads that mapped to BCR::ABL1 transcript in parental and TKI-resistant K562 cells treated with *Psp*Cas13b or imatinib (right). **(e)** RT-qPCR assays to measure the *Rfx*Cas13d-mediated knockdown of BCR-ABL1-p190 and eGFP using primers shown in Figure 2j, *N*=3. Data are normalized means and errors are SEM. Results are analysed by unpaired two-tailed Student’s t-test (95% confidence interval). *N* represents the number of biological replicates). Source data are provided as a Source data file.

This finding unveiled a surprising cleavage followed by potential RNA ligation that generates out-of-frame and translation-incompetent transcripts harbouring trimmed sequences indicated by increased non-canonical split reads (reads with no contiguous alignment to the reference) in the targeting condition (Figure 3d). Importantly, the frequency of split reads uniquely aligned to the BCR::ABL1 breakpoint region, where *Psp*Cas13b binds and cleaves, was significantly (*p*=0.024) higher (∼16%) compared to reads mapped to the entire transcript (∼3%) (Figure 3d, **Supplementary Fig. 2d**). When we used RT-qPCR primer sets complementary to various regions of the BCR::ABL1 mRNA, we confirmed this finding, as only the primers flanking the binding/cleavage site showed efficient RNA cleavage. In contrast, primers amplifying distal regions showed no significant RNA degradation (**Figure 3e**). Together, these data suggest that the RNA cleavage activity of *Psp*Cas13b is highly specific, but involves a previously unsuspected and unusual cleavage/religation mechanism that regenerates stable but misassembled RNA fragments.

### Personalised Design of *Psp*Cas13b Enables Potent Silencing of BCR::ABL1 RNA in Patient-Derived Cancer Cells

Next, we assessed whether our approach could specifically silence endogenous BCR::ABL1 fusion transcripts and suppress the growth of patient-derived cancer cell lines while leaving the wildtype BCR or ABL1 transcripts intact. K562 cells were originally derived from an adult patient with CML in blast crisis, and express the p210 variant of BCR::ABL1^29^. We personalized the design of PspCas13b to recognize the breakpoint sequence of p210 BCR::ABL1 variant. Using the IRES-eGFP reporter in HEK 293T cells, we confirmed the potent silencing efficiency (>95%) by crRNA0 (**Figure 1g**). We then tested whether crRNA0 could silence BCR::ABL1 mRNA in K562 cells. As K562 cells are refractory to conventional transfection, and lentiviral transduction of *Psp*Cas13b is largely ineffective^37^, we developed sequential nucleofection as a new approach to deliver *Psp*Cas13b and crRNA plasmids to K562 cells, followed by sorting of the *Psp*Cas13b-positive cells using BFP as a marker of uptake (**Supplementary Fig. 3a**). RT-qPCR analysis with primers designed to specifically detect BCR::ABL1 breakpoint, wildtype BCR and wildtype ABL1 confirmed efficient silencing of BCR::ABL1 mRNA, but no silencing of untranslocated BCR and ABL1 mRNA, demonstrating the high specificity of *Psp*Cas13b (**Figure 4a**). Consistent with RT-qPCR, western blot and FACS analysis demonstrated efficient silencing of BCR::ABL1 and the inhibition of STAT5 and ERK phosphorylation in the K562 cells, similar to that achieved with imatinib (**Figure 4b & 4c, Supplementary Fig. 3b & 3c)**. As K562 cells are highly dependent on BCR::ABL1 for proliferation and survival, we found that its silencing with *Psp*Cas13b greatly impaired cell proliferation over 96h, comparable to that achieved with imatinib (**Figure 4d & 4e**).

**Figure 4.**
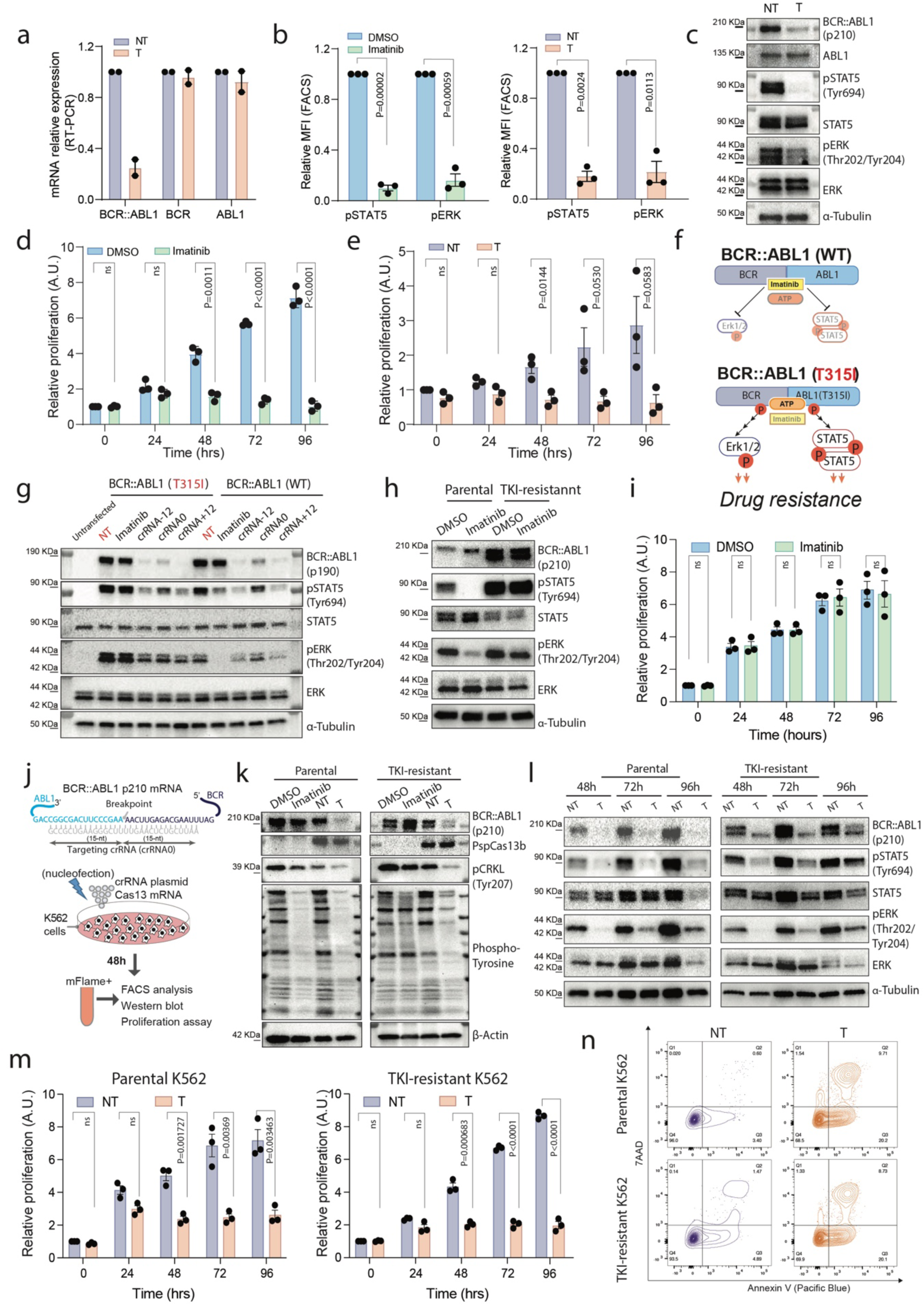
*Psp*Cas13b-mediated silencing of BCR::ABL1 in parental and TKI-resistant K562 cells suppresses downstream oncogenic signalling and cell proliferation. **(a)** RT-PCR assays to measure the expression levels of BCR::ABL1 p210, BCR and ABL1 mRNAs in K562 cells expressing non-targeting (NT) crRNA or crRNA0 targeting (T) BCR::ABL1 p210 mRNA, *N*=2. Data are normalized means and errors are SEM. **(b)** Intracellular cell flow cytometry measuring STAT5 and ERK phosphorylation levels in K562 cells following imatinib treatment or BCR::ABL1-p210 silencing with *Psp*Cas13b/T crRNA; *N*=3. Data are normalized mean fluorescence intensity (MFI) and errors are SEM; Results are analysed by unpaired two-tailed Student’s t-test (95% confidence interval). **(c)** Representative western blot analysis to examine the expression levels of BCR::ABL1-p210, untranslocated ABL1, and STAT5/ERK phosphorylation in K562 cells expressing NT or T crRNA; *N*=2 (See uncropped in the Source file). **(d-e)** Alarm blue assays to assess K562 cell proliferation following **(d)** imatinib treatment or **(e)** BCR::ABL1-p210 mRNA silencing with *Psp*Cas13b throughout a 96h time window (timepoint 0 is 24h post-transfection); *N*=3. Data are normalized means and errors are SEM; Results are analysed by unpaired Student’s T-test (95% confidence interval). **(f)** The schematic illustrates the imatinib-sensitivity or imatinib-resistance of wildtype (WT) BCR::ABL1 and T315I variants, respectively. **(g)** Representative western blot analysis to examine the suppression of T315I BCR::ABL1-p190 with *Psp*Cas13b. BCR::ABL1 expression and STAT5/ERK phosphorylation were examined in HEK 293T cells expressing WT or T315I BCR::ABL1 variants, *Psp*Cas13b, and either NT or T crRNAs (crRNA-12, crRNA0 and crRNA+12 previously assessed in Figure 1g) 24 h post-transfection. HEK 293T cells expressing BCR::ABL1 variants and *Psp*Cas13b were treated with 1µM imatinib for 4 hours as a positive control. HEK 293T cells were transfected with *Psp*Cas13b, NT and a control plasmid, which shows the baseline expression of pSTAT5 and pERK; *N* = 3 (See uncropped in the Source file). **(h)** Representative western blot analysis to confirm the imatinib-sensitivity or imatinib-resistance of parental and TKI-resistant K562 cells, respectively. *N* = 3 (See uncropped in the Source file). **(i)** Alamar blue assays to assess proliferation of TKI-resistant K562 cells following imatinib treatment throughout a 96h time window; *N*=3. Data are normalized means and errors are SEM; Results are analysed by unpaired Student’s T-test (95% confidence interval). **(j)** The schematic illustrates the experimental pipelines to assess *Psp*Cas13b*-*mediated silencing of BCR::ABL1-p210 in parental or TKI-resistant K562 cells using T crRNA (crRNA0). **(k)** Representative western blot analysis to examine the expression levels of *Psp*Cas13b BCR::ABL1-p210 and tyrosine phosphorylation in K562 cells expressing either NT or T crRNA 72 h post-transfection. K562 cells treated with imatinib or DMSO for 48h were used as controls; *N*=3 (See uncropped in the Source file). **(l)** Representative western blot analysis to examine the expression levels of BCR::ABL1-p210 and STAT5/ERK phosphorylation in K562 cells expressing either NT or T crRNA 48, 72 and 96 h post-transfection; *N*=3 (See uncropped in the Source file). **(m)** Alamar blue assays to assess proliferation of K562 cells following BCR::ABL1-p210 mRNA silencing with *Psp*Cas13 throughout a 96h time window (timepoint 0 is 24h post-transfection); *N*=3. Data are normalized means and errors are SEM; Results are analysed by unpaired Student’s T-test (95% confidence interval). **(n)** Representative FACS plots show K562 cells, parental (top) and TKI-resistant (bosom) expressing NT or T crRNA for 72 hours, then stained with Annexin V (Pacific Blue) and 7-ADD. Similar results were obtained in *N*=3. *N* is the number of independent biological replicates. Source data are provided as a Source data file.

Together, these data confirmed that *Psp*Cas13b could potently and specifically silence fusion drivers in patient-derived cell lines, shut down their oncogenic signalling, and impair cancer cell proliferation without silencing untranslocated transcripts.

**Supplementary Figure 3.**
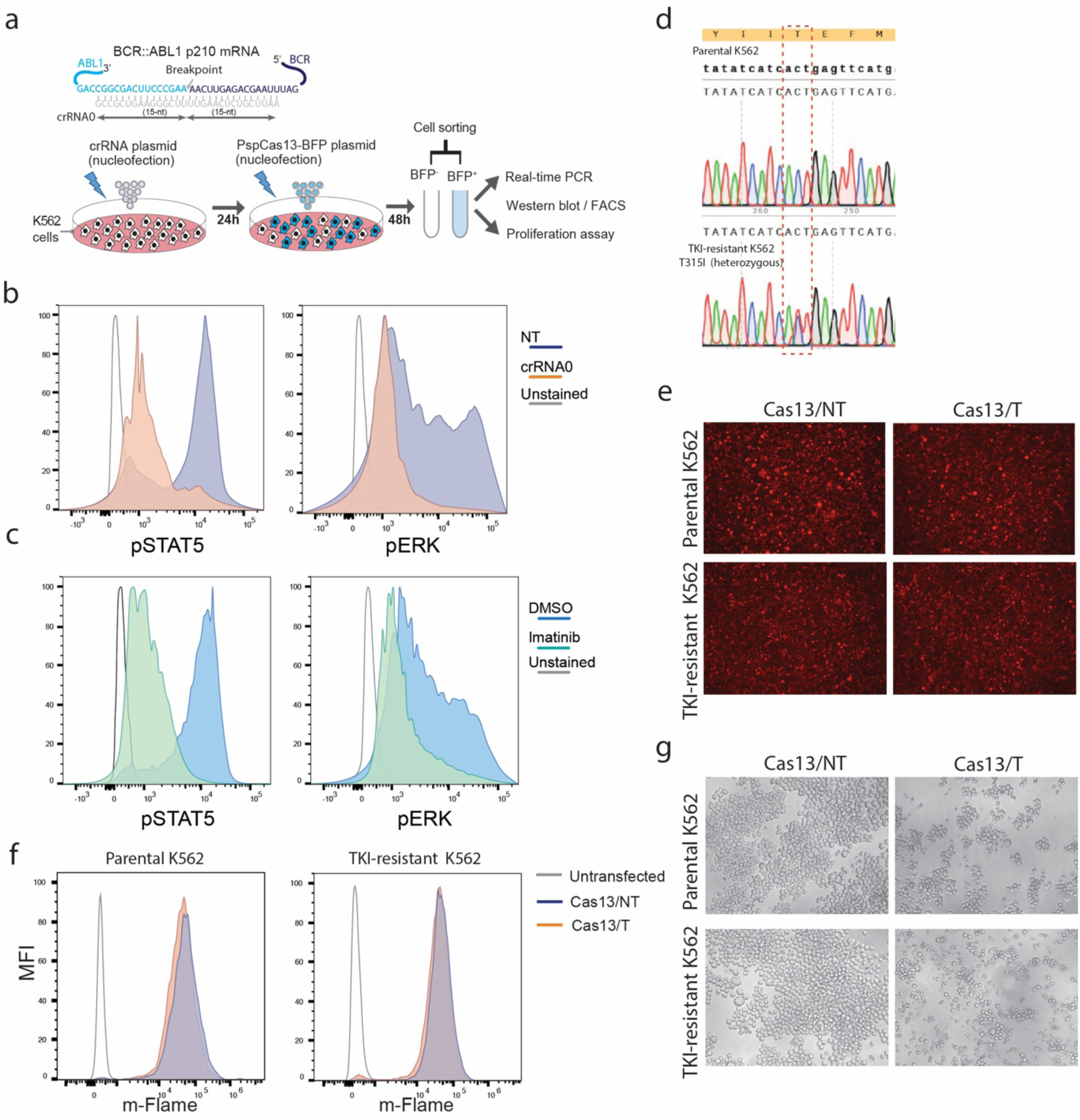
**(a)** The schematic illustrates the binding site of targeting (T) crRNA (crRNA0 in this case) to the breakpoint of BCR::ABL1-p210 mRNA, and experimental pipelines to assess *Psp*Cas13b*-*mediated silencing of BCR::ABL1 p210 in K562 cells using crRNA0. **(b-c)** Representative histograms of pSTAT5 and pERK staining in K562 cells expressing *Psp*Cas13/crRNA **(b)** or treated with imatinib **(c)**, *N*=3. **(d)** Sanger sequencing confirmed the heterozygous expression of T315I mutation of BCR::ABL1 in TKI-resistance K562 cells. **(e-f)** Representative fluorescence microscopy images **(e)** and histograms **(f)** show the expression of *Psp*Cas13b-mFlame in K562 cells expressing a non-targeting (NT) control crRNA versus a crRNA0 targeting (T) crRNA, *N* = 3. **(g)** Representative microscopy images show K562 cell density, parental (top) and TKI-resistant (bosom), at 96h time points post *Psp*Cas13b/crRNA delivery in cells expressing either NT or crRNA0 targeting (T) the BCR::ABL1-p210 mRNA. Scale bar = 300 μm. Similar results were obtained in *N*=3. Unprocessed representative images are provided in the Source Data file. *N* is the number of independent biological experiments. Source data are provided as a Source data file.

### *Psp*Cas13b Overcomes TKI-Resistance Driven by Secondary Mutations

Drug resistance acquired through secondary mutations remains a major challenge for all ABL1 kinase inhibitors currently approved to treat BCR::ABL1-driven leukemias^38^. For instance, the BCR::ABL1 kinase domain mutation Thr315Ile (T315I) confers resistance to ABL1 kinase inhibitors such as imatinib, nilotinib, or dasatinib and drives tumour relapse^31^ (**Figure 4f**). We hypothesised that unlike TKIs (e.g., imatinib), targeting the breakpoint of BCR::ABL1 transcript with *Psp*Cas13b should remain effective against both BCR::ABL1 and T315I BCR::ABL1 variants, as the single-nucleotide mutation that confers resistance is located outside of the targeted breakpoint sequence. To test this hypothesis, we compared the potency of imatinib and three *Psp*Cas13b crRNAs targeting the breakpoint of BCR::ABL1 in HEK 293T cells. As anticipated, imatinib efficiently inhibited the oncogenic signalling of BCR::ABL1 but not the T315I BCR::ABL1 variant (**Figure 4g**). Notably, all three *Psp*Cas13b crRNAs we tested largely inhibited the expression of both ancestral and T315I BCR::ABL1 proteins and their downstream oncogenic signalling, as evidenced by the inhibition of STAT5 and ERK phosphorylation. Consistent with previous data, crRNA-12 and crRNA+12 achieved the highest inhibitory effect due to higher silencing potency (**Figure 4g**). Next, we assessed whether our *Psp*Cas13b approach could suppress the oncogenic pathways in K562 cells that have evolved resistance to TKIs. This resistant cell line was generated by continuous, escalating dose exposure to dasatinib over >90 days until a resistant population was generated and maintained in 200 nM dasatinib^39^. Sanger sequencing confirmed that this cell line harboured a heterozygous point mutation (C>T) at the gatekeeper residue threonine-315 resulting in the substitution to isoleucine in this position (**Supplementary Fig. 3d**). We first confirmed that the parental K562 cells expressing ancestral BCR::ABL1 were sensitive to imatinib treatment as evidenced by the downregulation of pSTAT5 and pERK. The TKI-resistant K562 cells showed a significantly higher level of BCR-ABL1 protein compared to parental K562 cells (**Figure 4h**). STAT5 and ERK phosphorylation persisted in the TKI-resistant cells when treated with imatinib, and cell proliferation remained unaffected, consistent with imatinib resistance (**Figure 4h & 4i**). Next, we tested the ability of *Psp*Cas13b to silence T315I BCR::ABL1 in these TKI-resistant K562 cells. We optimised Cas13 delivery using nucleofection of *Psp*Cas13b-mFlame mRNA and crRNA plasmid (**Figure 4j**). This optimised nucleofection method achieved nearly 100% *Psp*Cas13b expression, as indicated by mFlame fluorescence, and resulted in minimal cell toxicity compared to the previous nucleofection protocol (**Supplementary Fig. 3e & 3f).** Efficient silencing of BCR::ABL1 by the targeting crRNA (crRNA0) was confirmed with western blot analysis, showing inhibition of global tyrosine phosphorylation, as detected by phospho-tyrosine (4G10) antibody, and reduced CRLK, STAT5 and ERK phosphorylation in both parental and TKI-resistant K562 cells (**Figure 4k & 4l**). Furthermore, silencing of BCR::ABL1 by *Psp*Cas13b remarkably inhibited cell proliferation and promoted apoptosis of both parental and TKI-resistant K562 cells over 96h (**Figure 4m & 4n, Supplementary Fig. 3g**).

Together, these data demonstrate that RNA targeting of the BCR::ABL1 breakpoint sequence can overcome TKI resistance commonly observed in relapsed leukemia.

### *Psp*Cas13b Drives Extensive Transcriptomic and Proteomic Remodeling in TKI-Resistant Cells

To further understand the downstream effect of BCR::ABL1 silencing by *Psp*Cas13b in parental and TKI-resistant K562 cells (T315I), we purified RNA from cells harvested 72h after *Psp*Cas13b-mFlame mRNA nucleofection with either a non-targeting or BCR::ABL1 targeting crRNA (crRNA0). As a control, parental and TKI-resistant K562 cells were treated with either imatinib or DMSO. We then performed a comparative RNA sequencing analysis under these conditions (**Figure 5a**). Consistent with observations in HEK 293T cells, *Psp*Cas13b efficiently cleaved both wild-type and T315I BCR::ABL1 near its binding site, as evidenced by fewer reads matching the breakpoint sequence and a significant increase in non-canonical split reads. This effect was not observed with a non-targeting crRNA or imatinib (**Supplementary Fig. 4a-4c**). *Psp*Cas13b-mediated silencing of BCR::ABL1 induced significant transcriptomic alterations in both parental and TKI-resistant cells. However, imatinib treatment only altered RNA expression in parental K562 cells, with no impact on TKI-resistant cells (**Figure 5b & 5c**). Parental and TKI-resistant cell lines treated with *Psp*Cas13b showed similar differential gene expression to that of parental cells treated with imatinib (**Figure 5d**). *Psp*Cas13b silencing of BCR::ABL1 in both parental and TKI-resistant cells resulted in 241 upregulated transcripts and 159 downregulated transcripts, which were also observed with BCR::ABL1 inhibition by imatinib in parental cells (**Supplementary Fig. 5a)**. These transcripts are involved in porphyrin metabolism and cancer signalling, including JAK-STAT, Rap1, PI3K-Akt, and MAPK pathways (**Supplementary Fig. 5b**). The most significantly upregulated common transcripts included PPP2R2C, ESPN, BEST3, SLC4A1, RP1-202O8.2, RLN3, CRB-55O6.4, and PHOSPHO1. Conversely, CD3D, BMP10, AC092566.1 and WNT7A were among the most significantly downregulated transcripts (**Supplementary Fig. 5c**). Gene ontology analysis of the top 60 upregulated and 60 downregulated transcripts revealed their involvement in positive regulation of megakaryocyte differentiation, myeloid cell differentiation, hemopoiesis, myeloid leukocyte activation, and regulation of stress response (**Supplementary Fig. 5d**).

**Figure 5.**
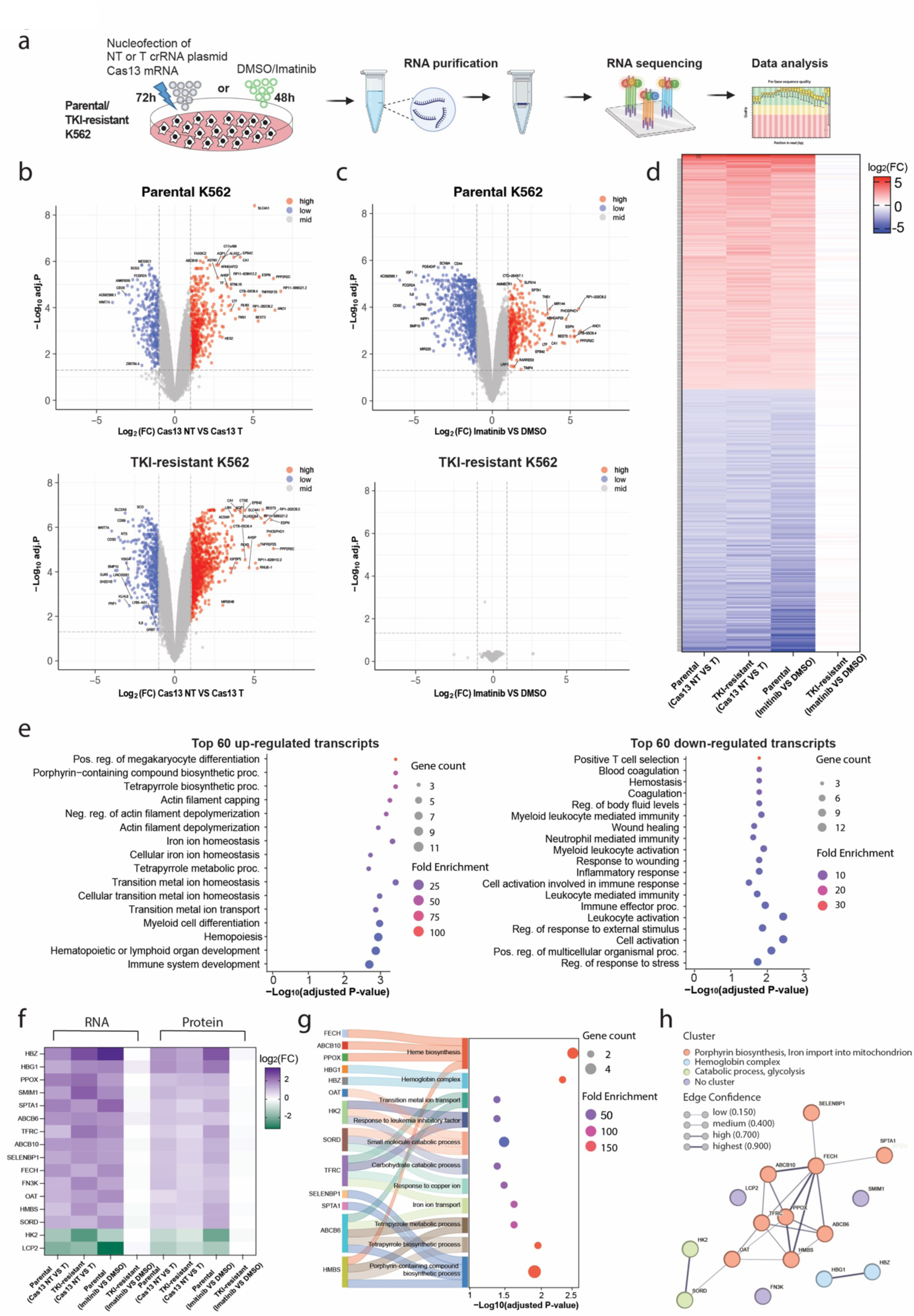
Transcriptomic and proteomic analysis reveals key molecular pathway alterations induced by *Psp*Cas13b-mediated BCR::ABL1 silencing in K562 cells. **(a)** Schematic of RNA sequencing and mass spectrometry assays to investigate the downstream effects of BCR::ABL1 silencing by *Psp*Cas13b in both parental and TKI-resistant K562 cells. The cells were nucleofected with *Psp*Cas13b-mFlame mRNA and either a targeting (T) or non-targeting (NT) crRNA plasmid. RNA and proteins were extracted 72 hours post-transfection. Cells treated with imatinib or DMSO for 48 hours were used as controls. **(b, c)** Volcano plots of RNA-sequencing analysis of the transcriptome of parental (top) and T315I (bosom) K562 cells expressing (b) *Psp*Cas13b or **(c)** treated with imatinib. Each data point represents a transcript, and transcripts with log_2_FC > 1 (upregulation, shown in red) or log_2_FC < -1 (downregulation, shown in blue), and a Benjamini-Hochberg adjusted p-value < 0.05 are considered significant differentially expressed transcripts. *N*=3. **(d)** Heatmap for 364 common transcripts differentially expressed in *Psp*Cas13b treated cell lines (parental or TKI-resistant) and imatinib-treated parental cells. **(e)** Gene ontology analysis for differentially expressed proteins shown in mass spectrometry data in **Supplementary Fig. 6b and 6c**. **(f)** Gene ontology analysis for the 32 common differentially expressed proteins (25 upregulated and 7 downregulated) in *Psp*Cas13b/T crRNA expressing K562 (parental or TKI-resistant) cells and imatinib-treated parental cells shown in the Venn diagram **Supplementary Fig. 6d**. The gradient colour scheme represents the averaged log_2_FC of each protein’s expression in *Psp*Cas13b/T crRNA expressing K562 (parental or TKI-resistant) or imatinib-treated parental cells relative to their corresponding control. **(g)** Heatmap for 16 genes differentially expressed in cell lines (parental or TKI-resistant) treated with *Psp*Cas13b and parental cells treated with imatinib compared to their respective control groups. These genes were confidently detected by both RNA sequencing and mass spectrometry across all four comparison groups. **(h)** Gene ontology analysis for the 16 genes shown in the heatmap **f**. LCP2, SMIM1, and FN3K are not shown, as a minimum of two genes in each pathway is required. **(i)** STRING Protein-protein interaction networks functional enrichment analysis for the 16 genes shown in the heatmap **g**. *N* represents the number of biological replicates. Source data are provided as a Source data file.

To gain deeper insights into the remodelling of proteome due to *Psp*Cas13b silencing of BCR::ABL1, we performed mass spectrometry analysis on parental and TKI-resistance K562 cells (**Figure 5a).** Consistent with RNA sequencing, *Psp*Cas13b effectively downregulated BCR::ABL1 in both cell types, resulting in significant proteomic alterations (**Supplementary Fig. 6a-6b**). In contrast, imatinib treatment induced a drastic alteration in the proteome of parental K562 cells, but had a minimal impact on TKI-resistant K562 cells, consistent with the anticipated TKI-resistance phenotype (**Supplementary Fig. 6c**). Next, we investigated whether BCR::ABL1 inhibition using either *Psp*Cas13b or imatinib in parental and TKI-resistant K562 cells would result in similar proteomic signatures. In TKI-resistant K562 cells treated with *Psp*Cas13b, 47 proteins were upregulated and 29 were downregulated, with 37 (78%) and 16 (55%) of these overlapped with proteins differentially expressed in parental cells treated with either imatinib or *Psp*Cas13b. (**Supplementary Fig. 6d**).

**Supplementary Figure 4.**
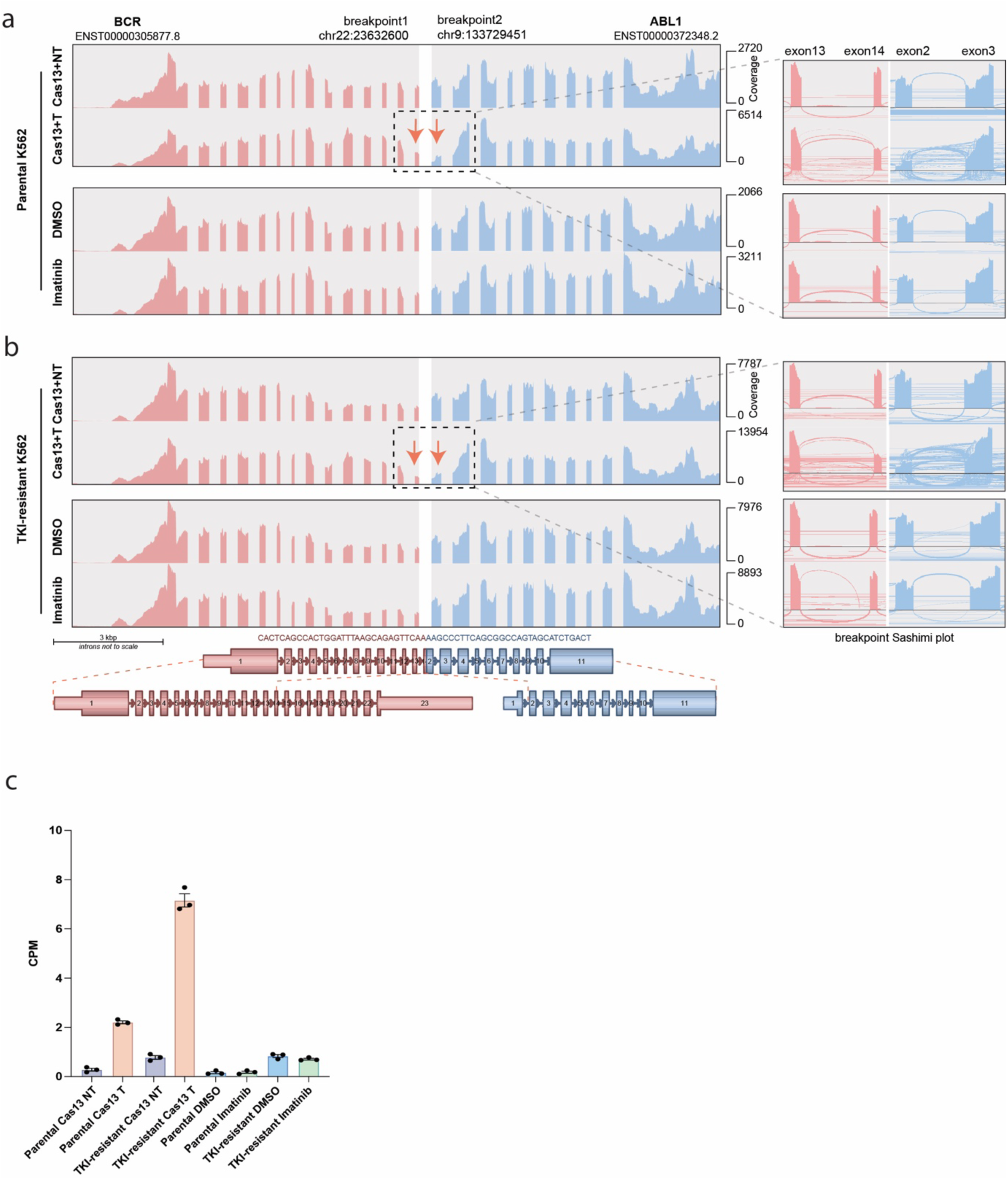
**(a-b)** Representative Arriba fusion plots (right) and Sashimi plots showing the read alignments against the BCR-ABL1 transcript in parental **(a)** and TKI-resistant **(b)** K562 cells expressing *Psp*Cas13b with either a non-targeting (NT) or a targeting (T) crRNA, or treated with either imatinib or DMSO. The breakpoint region is highlighted with a dashline box and the approximate cleavage site is indicated with a red arrow. **(c)** Counts per million (CPM) of non-canonical split reads that exclusively mapped to BCR::ABL1 breakpoint (filter criteria shown in the method section) in parental and TKI-resistant K562 cells treated with *Psp*Cas13b or imatinib. *N* represents the number of biological replicates). Source data are provided as a Source data file.

**Supplementary Figure 5.**
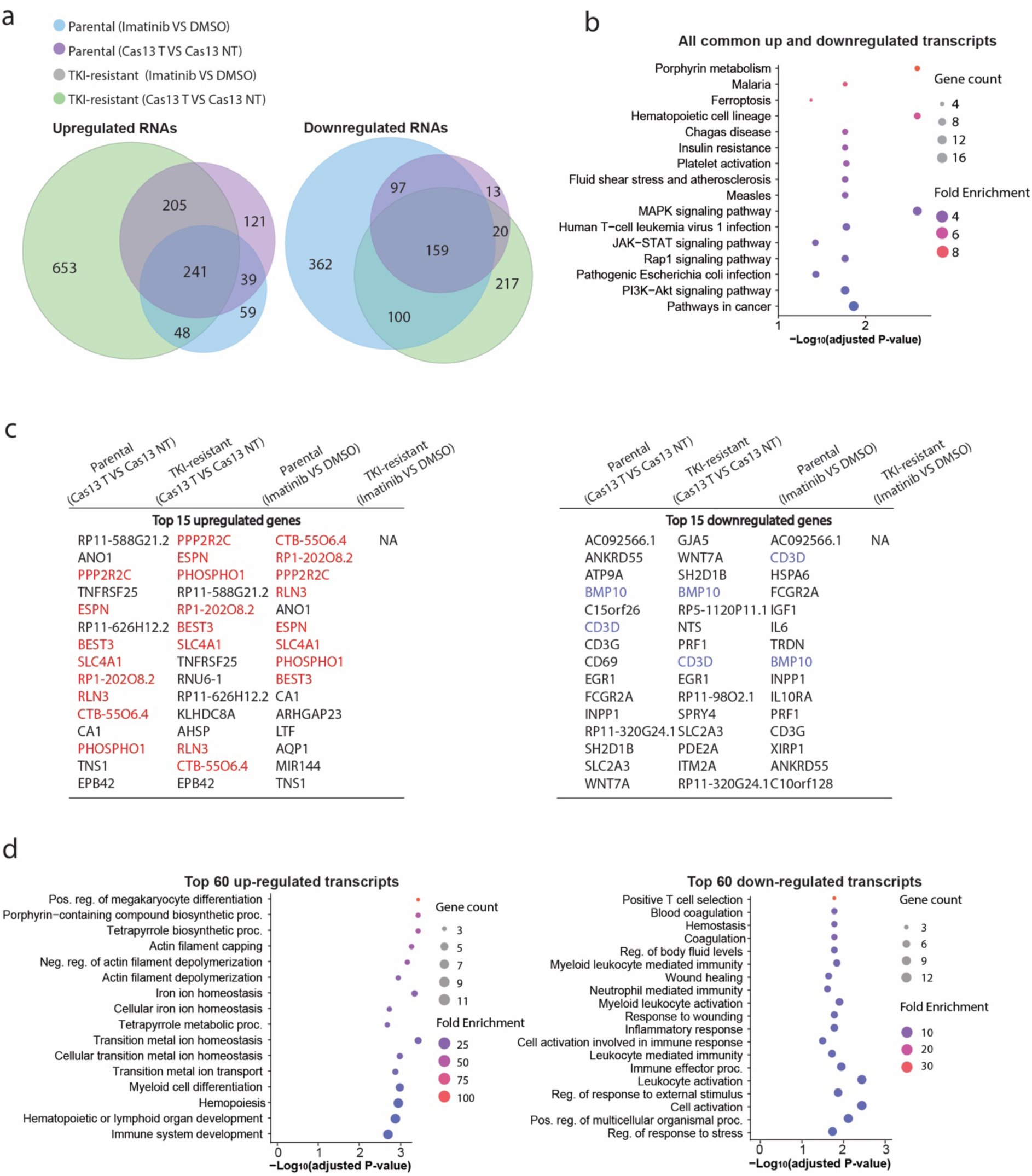
**(a)** Venn diagrams showing the number of overlapping and unique transcripts differentially expressed (defined by a log_2_ fold change >1 or <-1, with an adjusted p-value < 0.05) in RNA-sequencing data from the imatinib or *Psp*Cas13b treated K562 cells relative to the corresponding control K562 cells. **(b)** Gene ontology analysis of the common 400 transcripts differentially expressed in *Psp*Cas13b treated cell lines (parental or TKI-resistant) and imatinib-treated parental cells shown in the Venn diagrams **a**. **(c)** The list of top 15 upregulated and downregulated proteins from the RNA sequencing analysis in parental and TKI-resistant K562 cells treated with *Psp*Cas13b or imatinib. **(d)** Gene ontology analysis for the top 60 up-regulated and 60 downregulated transcripts shown in the heatmap Figure 5d.

The significant overlap in the protein signatures mediated by BCR::ABL1 inhibition with imatinib in parental cells, and *Psp*Cas13b in both parental and TKI-resistant cells was further highlighted by the heatmap analysis. However, the proteome of TKI-resistant cells treated with imatinib did not show any substantial overlap with the other three conditions (**Supplementary Fig. 6e**). Pathway enrichment and protein network analyses of the most significantly upregulated proteins induced by both *Psp*Cas13b and imatinib treatments revealed their key roles in haemoglobin complex formation (HBG1, HBG2, HBZ, HBE1, HBA1, HBA2, HBD), heme biosynthesis (PPOX, HMBS, FECH), erythropoiesis (TFRC, SPTA1, SMIM1), mitochondrial (ABCB10, ABCB6, OAT, HAGH), and positive regulation of cell death (PDCD4). Conversely, the most significantly downregulated proteins were involved in lipid metabolism (FADS1, CYP51A1, HMGCS1, IDI1), glucose metabolism (HK2), and cell signalling (LCP2, CDKAL1) (**Figure 5e-5f, Supplementary Fig. 6f, Supplementary Fig. 7a-7d)**.

**Supplementary Figure 6.**
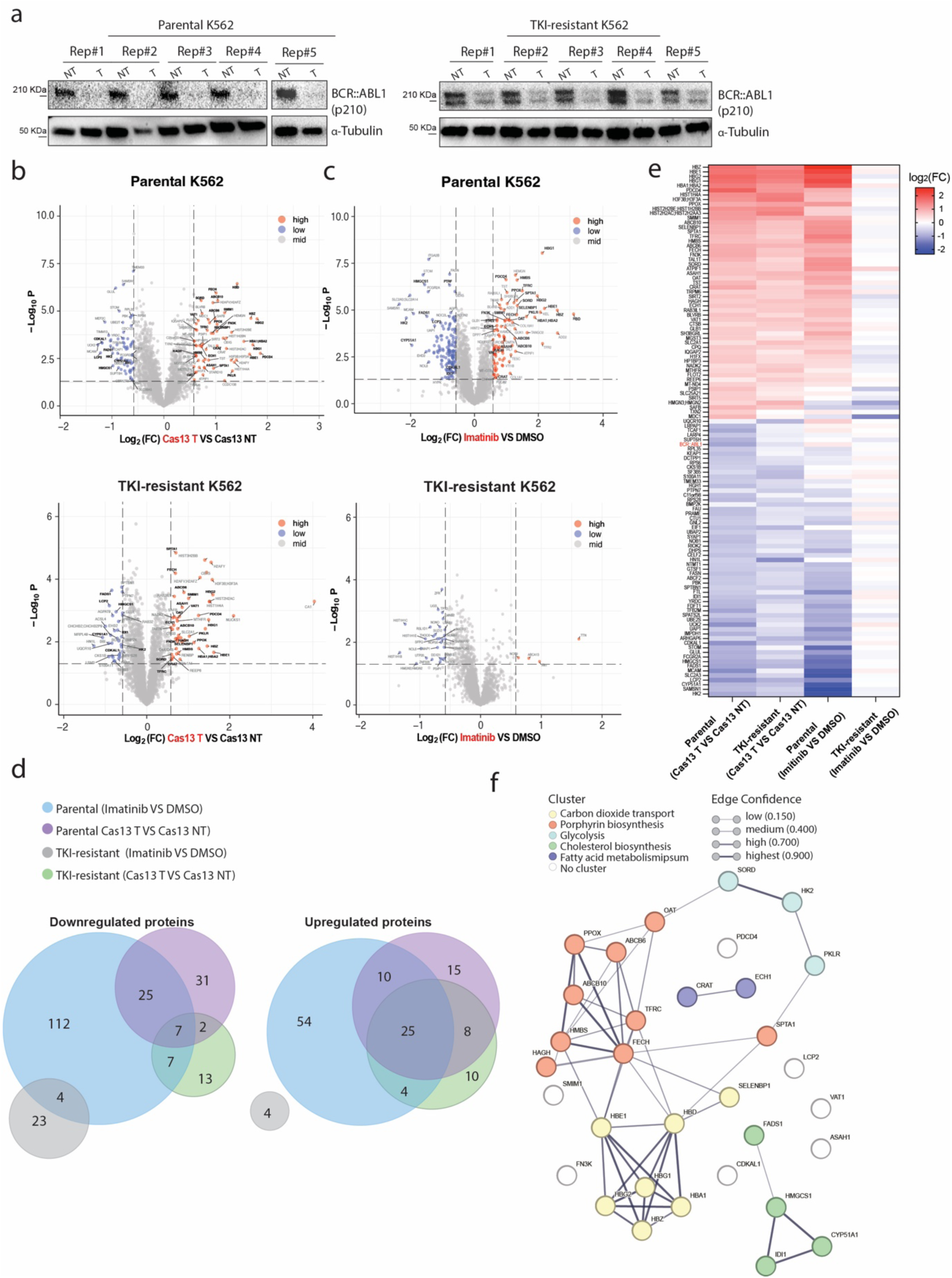
**(a)** Western Blot validation of BCR::ABL1 expression in TKI-resistant and parental K562 cells expressing *Psp*Cas13b with either a non-targeting (NT) crRNA or a targeting (T) crRNA, using the same lysates as those used for mass spectrometry analyses, *N* = 5 (See uncropped blots in the Source file). **(b, c)** Volcano plots of mass spectrometry analysis of the proteome of parental (top) and T315I (bosom) K562 cells expressing **(b)** *Psp*Cas13b or **(c)** treated with imatinib. Each data point represents a protein, and proteins with log_2_FC > 0.5 (upregulation, shown in red) or log_2_FC < -0.5 (downregulation, shown blue), and p-value < 0.05 are considered significant differentially expressed proteins. Common up or down-regulated proteins of *Psp*Cas13b-treated cells and imatinib-treated TKI-resistant K562 cells are labelled in bold in the plot. *N*=5. **(d)** Venn diagrams showing the number of overlapping and unique proteins differentially expressed (calculated as log_2_ fold change >1 or <-1, with an adjusted p-value <0.05) in mass spectrometry data from the imatinib or *Psp*Cas13b treated K562 cells relative to the corresponding control K562 cells. **(e)** Heatmap for 115 proteins differentially expressed at log_2_FC >0.5 or <−0.5 in at least one of the *Psp*Cas13b treated cell lines (parental or TKI-resistant). The colour scheme represents the log_2_FC of each protein’s expression in Imatinib or *Psp*Cas13b treated K562 cells relative to their corresponding control. **(f)** STRING Protein-protein interaction networks functional enrichment analysis for the 32 common up (25) and down-regulated (7) proteins shown in the Venn diagrams in **d** and the chord diagram in Figure 5f. *N* is the number of independent biological experiments. Source data are provided as a Source data file.

We performed further analysis to integrate transcriptomic and proteomic data and assess the correlation between transcriptomic and proteomic alterations. Approximately 2,800 proteins (90%) detected by mass spectrometry in each experimental condition have corresponding transcripts in RNA-seq data. Of these, 2,516 genes were consistently matched across all four comparison groups with high confidence at both the transcript and protein levels, revealing a positive correlation between RNA expression and their corresponding proteins in parental cells treated with *Psp*Cas13b or imatinib, as well as in TKI-resistant cells treated with *Psp*Cas13b (**Supplementary Fig. 8**). Importantly, 16 genes exhibited significant differential expression with high confidence at both the RNA and protein levels in parental and TKI-resistant cells treated with *Psp*Cas13b, as well as in parental cells treated with imatinib (**Figure 5g**). Gene ontology and protein-protein interaction analyses of these altered 16 candidates revealed their implication in the regulation of porphyrin biosynthesis (PPOX, SPTA1, ABCB6, TFRC, ABCB10, SELENBP1, FECH, OAT, and HMBS), haemoglobin complex assembly (HBZ and HBG1), and glycolysis (HK2 and SORD) (**Figure 5h & 5i**).

**Supplementary Figure 7.**
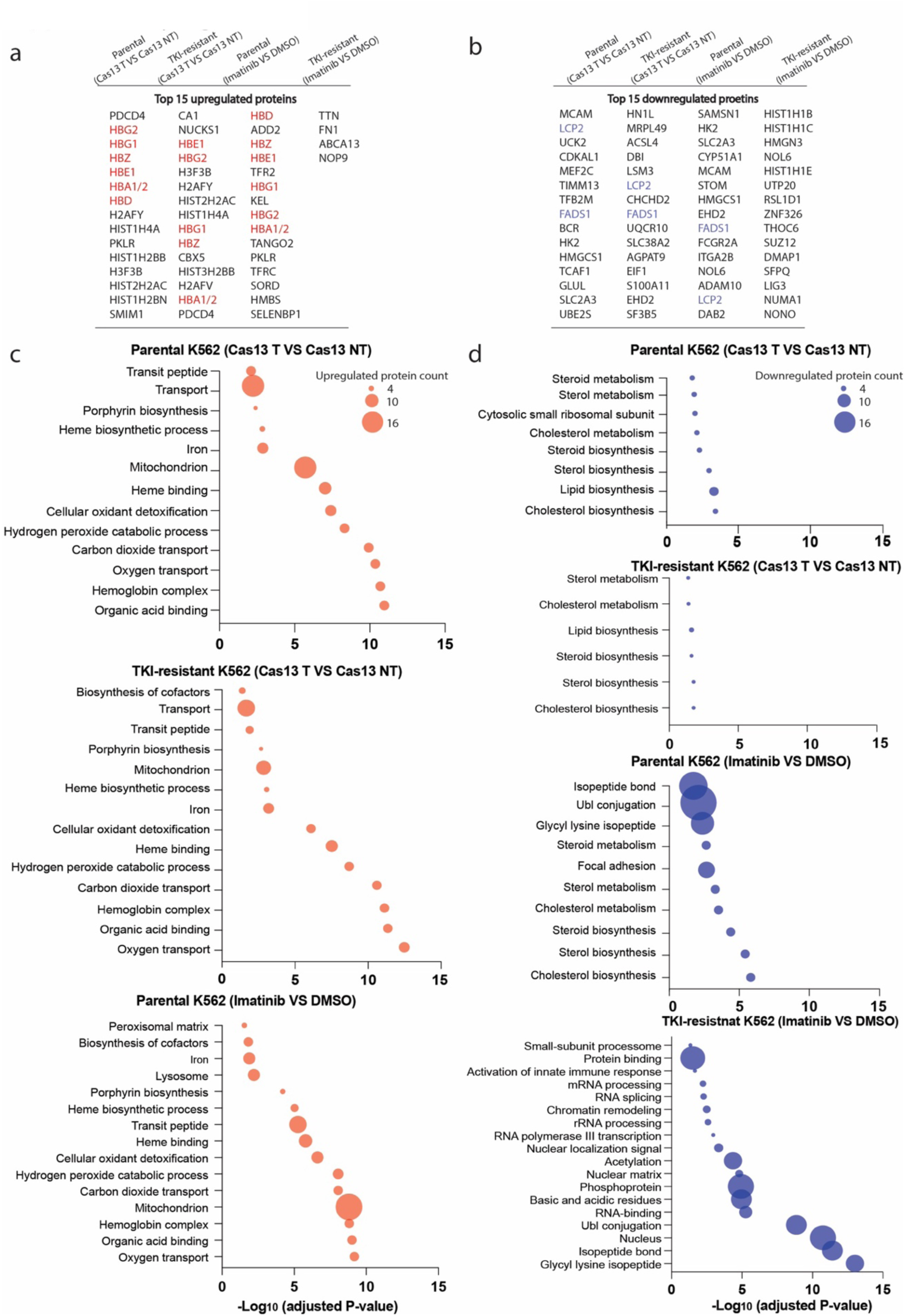
**(a-b)** The list of top 15 upregulated **(a)** and downregulated proteins **(b)** from the proteomic analysis in parental and TKI-resistant K562 cells treated with *Psp*Cas13b or imatinib. **(c-d)** Gene ontology analyses of upregulated **(c)** and downregulated **(d)** proteins in parental and TKI-resistant K562 Cells treated with *Psp*Cas13b or imatinib. Source data are provided as a Source data file.

**Supplementary Figure 8.**
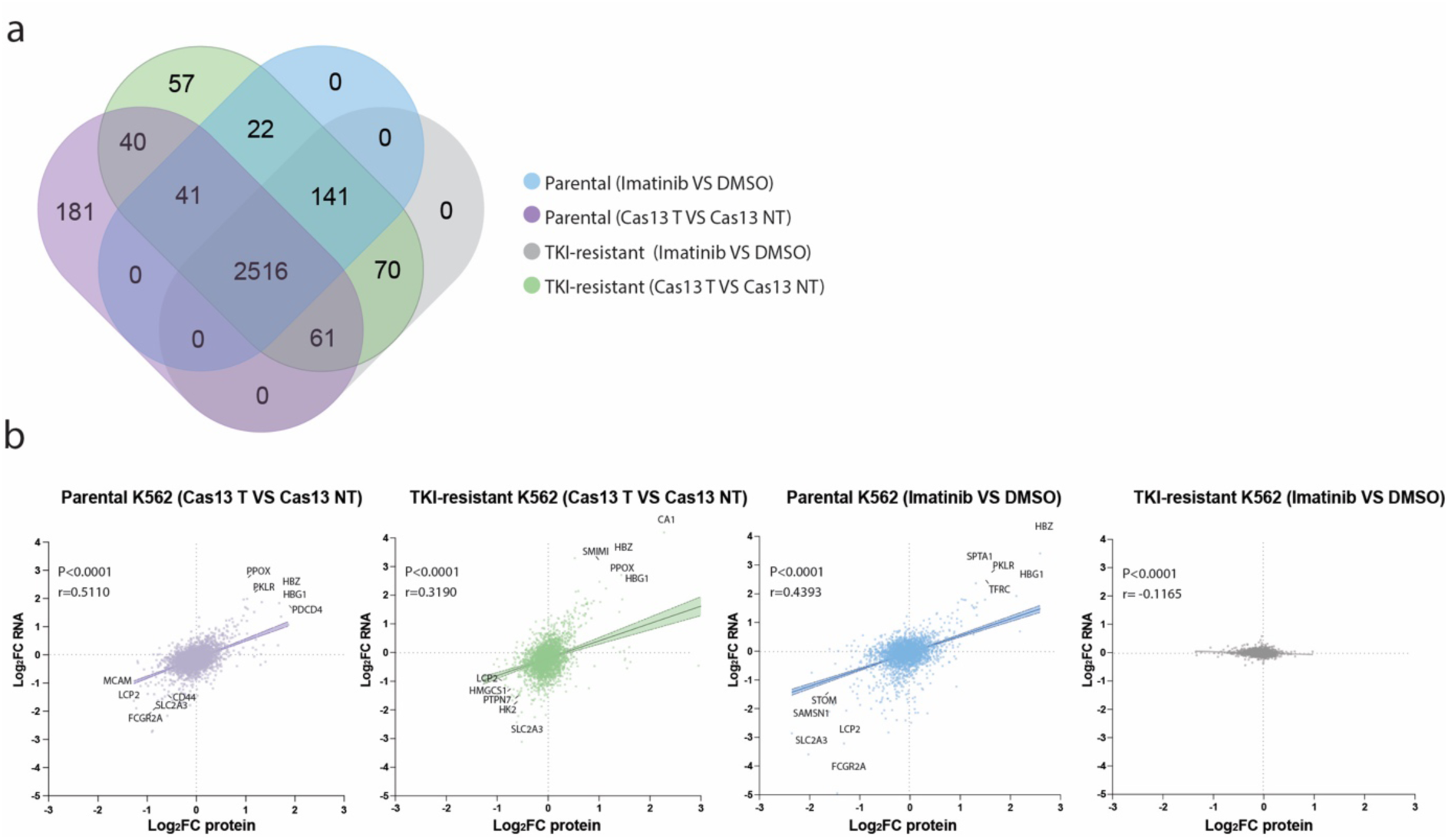
Protein-level alterations induced by *Psp*Cas13b-mediated BCR::ABL1 silencing in K562 cells are consistent with transcript-level observations. **(a)** Venn diagrams display the number of overlapping and unique genes confidently detected by both RNA sequencing and mass spectrometry in K562 cells treated with imatinib or *Psp*Cas13b, as well as in the corresponding control cells. **(b)** Relationship between RNA and protein log_2_ fold change (FC) levels of genes measured by RNA sequencing and mass spectrometry in K562 cells treated with imatinib or *Psp*Cas13b, relative to their corresponding control cells. Pearson correlation analyses were performed. *r* (correlation coefficient) and *p-*value (95% confidence interval) are indicated in each graph. Results are analysed by unpaired two-tailed Student’s t-test. Error bars are SEM. Source data are provided as a Source data file

Together, these data demonstrate that the profound remodelling of the transcriptome and proteome of TKI-resistant cells through T315I BCR::ABL1 transcript targeting with *Psp*Cas13b forces these cells into erythroid differentiation and apoptosis, profoundly altering their drug resistance and tumorigenic properties.

### Efficient Silencing of Emerging Drug-Resistant Mutations

The next-generation of BCR::ABL1 inhibitors such as ponatinib (ATP-binding inhibitor) and asciminib (allosteric inhibitor) have been developed to inhibit the most frequent drug-resistant mutations including T315I^40,41^. However, non-responsive BCR::ABL1 isoforms^42^ and newly emerging mutations^43^ have been reported to confer resistance to these advanced drugs. For example, mutations such as C464W, A337V and P465S have been identified as conferring resistance to asciminib^43^. We reasoned that PspCas13b could eliminate ascitinib-resistant BCR-ABL1 transcripts, irrespective of the type or position of the mutation. To test this, we generated asciminib-resistant BCR::ABL1 mutants (C464W and T315I-C464W) and showed that *Psp*Cas13b targeting the breakpoint of BCR::ABL1, effectively silenced these drug-resistant mutants. In contrast, ascitinib suppressed the wildtype BCR-ABL1, but failed to inhibit C464W and T315I-C464W mutants (**Supplementary Fig. 9a**). This data indicates that targeting fusion transcripts with PspCas13b remains effective against emerging drug-resistant mutations.

**Supplementary Figure 9.**
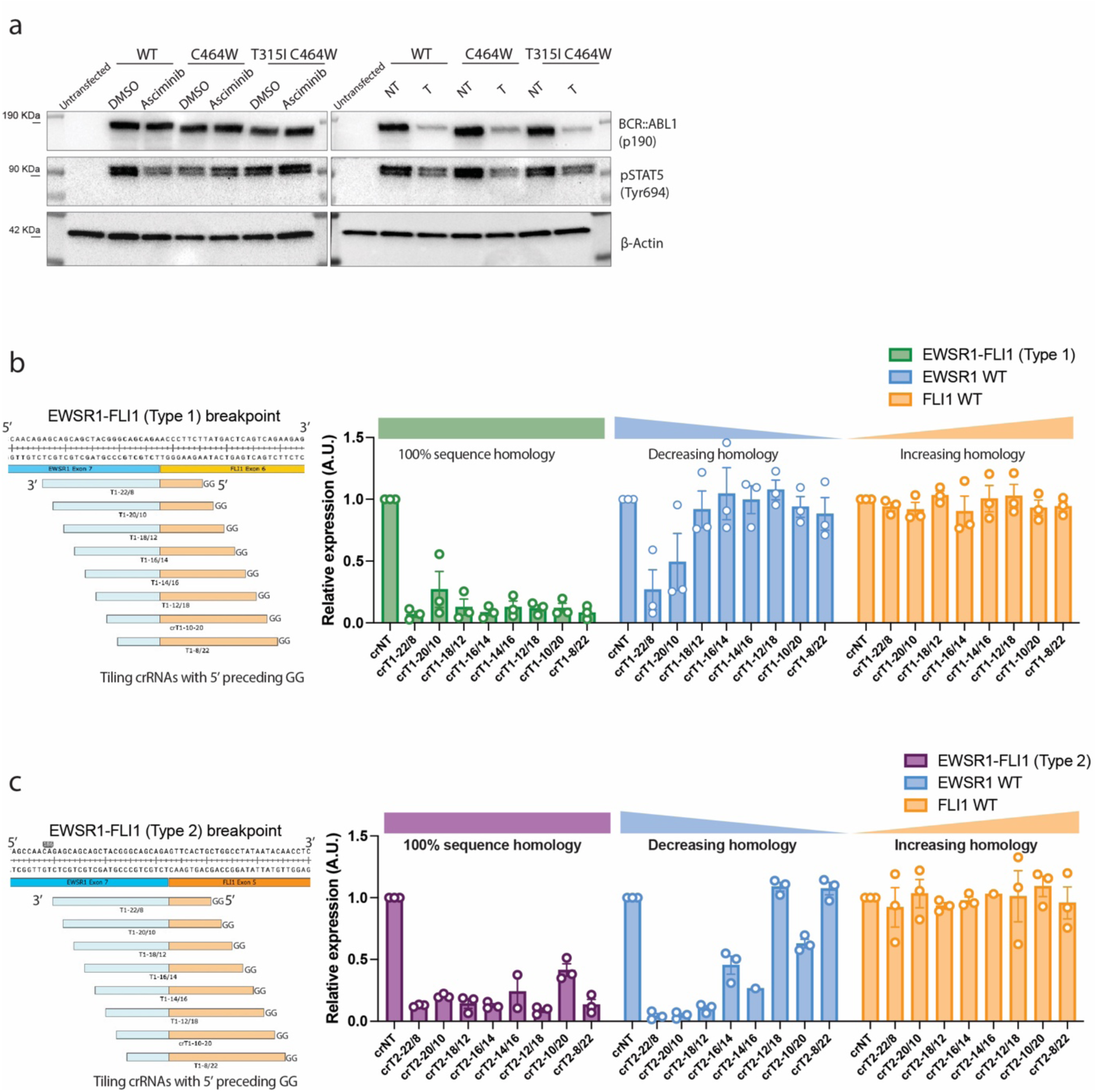
The targeting of the breakpoint of EWSR1-FLI can efficiently discriminate between translocated tumour RNAs and wildtype variants. **(a)** Representative western blot analysis to examine the suppression of C464W and T315I-C464W BCR::ABL1-p190 variants with *Psp*Cas13b. BCR::ABL1 expression and STAT5 phosphorylation were examined in HEK 293T cells expressing WT, C464W or T315I-C464W BCR::ABL1 variants, *Psp*Cas13b, and either NT or T crRNAs (crRNA0 previously assessed in Figure 1g) 24 h post-transfection. HEK 293T cells expressing BCR::ABL1 variants and *Psp*Cas13b were treated with 1µM asciminib for 4 hours as a positive control. Untransfected HEK 293T cells show the baseline expression of pSTAT5; *N* = 3 (See uncropped WB in the Source file). EGFP reporter assay to assess the silencing efficiency of tiled *Psp*Cas13b crRNAs incorporated with a 5’ GG motif with 2-nucleotide resolution targeting the breakpoint region of fusion transcripts EWSR1::FLI type 1 (EWSR1 Exon 7 to FLI1 Exon 6 translocation) **(b)** and type 2 (EWSR1 Exon 7 to FLI1 Exon 5 translocation) **(c)**. Data points in the graphs are averages of mean fluorescence from 4 representative fields of view per condition imaged. The data are represented in arbitrary units (A.U.). Errors are SEM.

### Personalised Design of *Psp*Cas13b Enables Potent Silencing of Various Fusion Transcripts

Next, we investigated whether the design strategy described above could enable us to predict crRNAs that would efficiently target other undruggable fusion transcripts. For instance, RUNX1::RUNX1T1 is an oncogenic fusion gene that recruits chromatin corepressors to block cell differentiation in AML^44^, while the oncogenic fusion NPM-ALK encodes a constitutively active tyrosine kinase that results in anaplastic large cell lymphoma^45^. Similarly, the SS18-SSX1/2 fusion gene is pivotal in the advancement of synovial sarcoma^46^. We generated five crRNAs that we predicted to efficiently target the breakpoint of each fusion transcript and tested their silencing efficiencies^5^. Using ectopic expression of gene fusions in HEK 293T cells, we demonstrated efficient silencing of the RUNX1::RUNX1T1, NPM::ALK, and SS18::SSX1/2 fusion transcripts with the personalised design of *Psp*Cas13b (**Figure 6a-6c)**. In addition, we demonstrated efficient and selective silencing of EWSR1::FLI1 fusion transcript, a well-established driver of Ewing sarcoma^47^ (**Supplementary Fig. 9c & 9d**).

**Figure 6.**
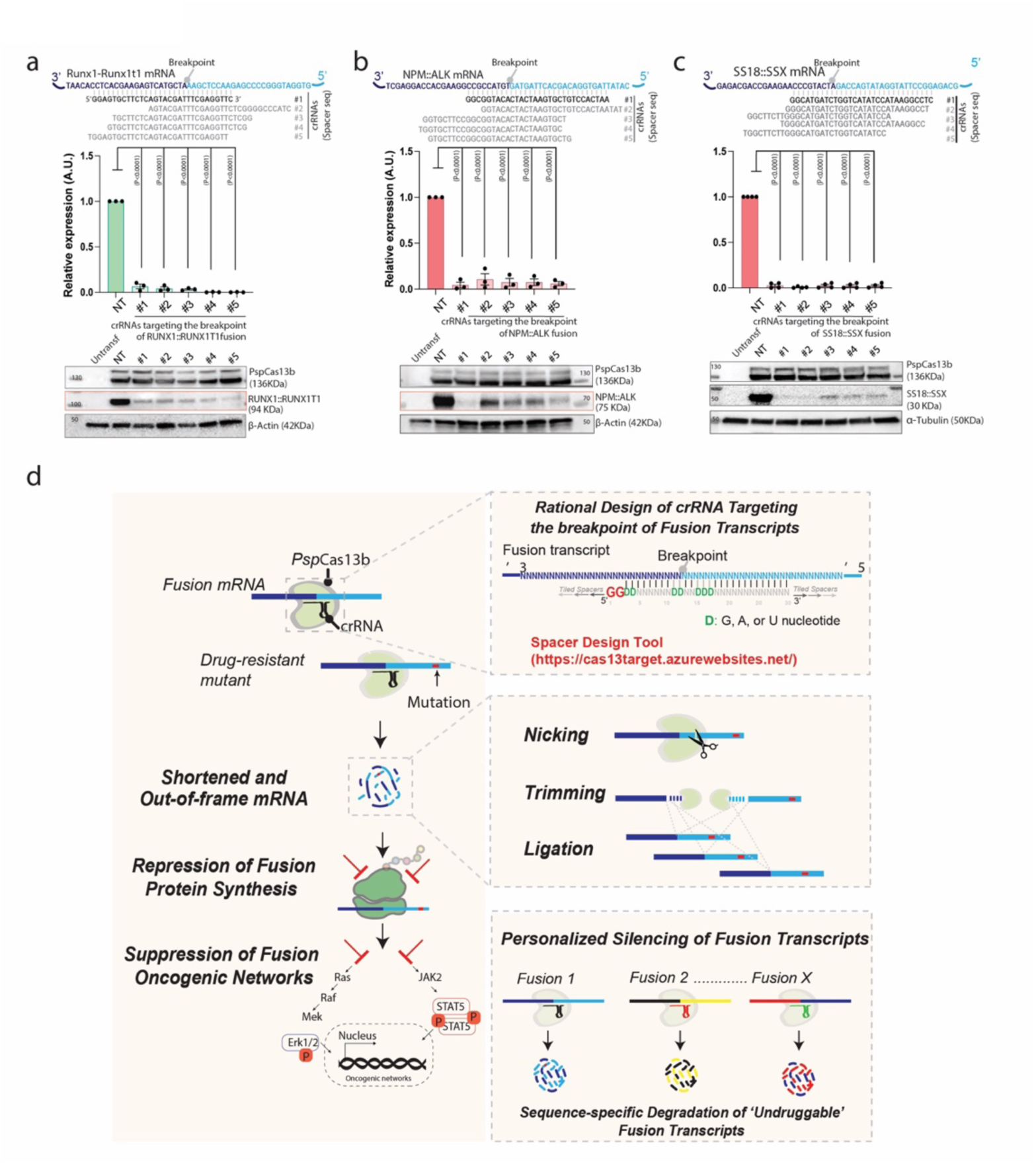
*Psp*Cas13b-mediated potent silencing of fusion transcripts in various cancer types. **(a-c)** Schematic (top) of predicted *Psp*Cas13b crRNAs targeting the breakpoint sequence of **(a)** RUNX1-RUNX1T1, **(b)** NPM-ALK and **(c)** SS18-SSX1/2 mRNAs, respectively. Quantifications (middle) of the silencing efficiency of predicted potent crRNAs targeting **(a)** RUNX1-RUNX1T1, **(b)** NPM-ALK and **(c)** SS18-SSX1/2 fusion transcripts in HEK 293T cells. Data points in the graph are averages of normalised mean fluorescence from 4 representative fields of view per experiment imaged in *N* = 3. The data are represented in arbitrary units (A.U.). Errors are SEM and *p*-values of the one-way ANOVA test are indicated (95% confidence interval). Representative western blot (bosom) analysis to examine the expression level of **(a)** RUNX1-RUNX1T1, **(b)** NPM-ALK and **(c)** SS18-SSX1/2 in HEK 293T cells; *N*=3 (See uncropped blots in the Source file). **(d)** Design principles for potent, specific, and personalised silencing of various gene fusion mRNAs with programmable *Psp*Cas13b nuclease. The lel panel shows the rational design of crRNAs targeting the breakpoint of gene fusion transcripts, which enables efficient and specific silencing of these oncogenic drivers, as well as drug-resistant fusion mutants. *Psp*Cas13b cleavage selectively trimmed a defined region near its binding site, leaving distal regions intact. The cleaved strands underwent RNA ligation, generating out-of-frame sequences that disrupted BCR::ABL1 translation. This targeting of the breakpoint strategy avoids the off-targeting of untranslocated gene partners that are expressed in normal tissues. The top right panel summarises spacer nucleotide positions that are important for the rational design of potent crRNA, which are integrated into our *in-silico* tool (hsps://cas13target.azurewebsites.net/)^5^. The bosom right panel shows the versatility and therapeutic potential of *Psp*Cas13b to silence various ‘*undruggable*’ gene fusion transcripts and their oncogenic networks in a personalised manner through precise targeting of fusion’s breakpoint sequence that is exclusively expressed in tumour cells. *N* is the number of independent biological experiments. Source data are provided as a Source data file.

Together, these data indicates that *Psp*Cas13b is a versatile platform capable of silencing a variety of gene fusions in a personalised manner.

## DISCUSSION

Over 3,297 unique fusion transcripts have been described across multiple cancer types^48^, most of which remain ‘*undruggable*’. In this proof-of-concept study, we demonstrate that personalized design of *Psp*Cas13b can efficiently recognise and silence nine different oncogenic fusion transcripts. To benchmark *Psp*Cas13b efficacy and specificity against clinically approved drugs, we focused this investigation on targeting BCR::ABL1 fusion, a well-established driver of leukemias^33,49–52^, which revealed key properties of fusion transcripts silencing with *Psp*Cas13b.

### High specificity

Although TKIs targeting BCR::ABL1 oncoprotein exhibit remarkable efficacy in controlling the progression of BCR::ABL1-driven leukemias, the use of TKIs in treating BCR::ABL1-positive leukemias is frequently associated with various degrees of toxicity due to the non-specific inhibition of ABL1 and ABL1-like kinases in normal tissues^53^. We showed that custom-designed *Psp*Cas13b targeting the breakpoint of BCR::ABL1 transcript exhibits high efficacy and exquisite specificity due to its low mismatch tolerance^5^ (**Figure 2**). *Psp*Cas13b silenced BCR::ABL1 transcripts and downregulated their oncogenic signalling pathways without off-targeting untranslocated BCR or ABL1 transcripts or other endogenous transcripts (**Figure 3**). This high specificity indicates that *Psp*Cas13b is highly unlikely to degrade untranslocated normal mRNAs expressed in non-cancerous cells despite their partial sequence homology with the fusion transcript.

### Unique cleavage pattern

RNA sequencing did not show a significant reduction in overall BCR::ABL1 mRNA levels, but closer analysis revealed selective degradation of a ∼500-nt long region near the *Psp*Cas13b cleavage site (**Figure 3, Supplementary Fig. 4**). In fact, RT-qPCR confirmed efficient cleavage only near the binding site (**Figure 3)**. This cleavage pattern was surprising, and indicated that the target RNA was not fully degraded, yet protein expression was greatly compromised (**Figure 1,2 & 4**). The frequency of non-canonical split reads was significantly higher around the cleavage site (∼16%) than the full transcript (∼3%) (**Supplementary Figure 2**), supporting the idea of targeted cleavage at the breakpoint followed by RNA ligation. The observed RNA nicking and ligation is unexpected, as *Psp*Cas13b is localised in the cytoplasm, while conventional RNA ligation in human cells during splicing occurs in the nucleus^54^. However, previous studies have shown that the cytoplasmic RNA ligase RTCB catalyses non-conventional, cytoplasmic splicing of the XBP1 mRNA during the unfolded protein response (UPR)^55,56^. The molecular mechanisms underlying the generation of these non-conventional trimmed transcripts following *Psp*Cas13b cleavage remain unclear and warrant further investigation (**Figure 6d**).

### Resilience to secondary mutations that drive drug resistance

In addition to drug toxicity, acquired resistance through secondary mutations remains a major challenge in the management of multiple cancers. For example, the BCR::ABL1 mutations T315I and C464W confer resistance to ABL1 TKIs, including imatinib and asciminib, and often drives tumour progression in patients^31^. Unlike imatinib, *Psp*Cas13b-mediated BCR::ABL1 silencing significantly inhibited cell proliferation and induced apoptosis in both parental and TKI-resistant cells (**Figure 4**). Although we only demonstrated that *Psp*Cas13b can overcome drug resistance against the T315I and C464W mutations, direct cleavage of the BCR-ABL1 transcript at the breakpoint is anticipated to suppress a wide spectrum of mutations and protein variants resistant to small inhibitory molecules. Additionally, *Psp*Cas13b can tolerate 3 to 4 nucleotide mismatches with its target (**Figure 2**)^5^, providing resilience to potential mutations if they occur within its 30-nucleotide binding site at the breakpoint sequence.

Transcriptomic and proteomic profiling of BCR::ABL1 silencing by *Psp*Cas13b offers insights into the potential of this approach as a therapeutic strategy for fusion gene-driven cancers, particularly in the context of drug resistance. Our data revealed a significant overlap in protein and transcript signatures between imatinib-treated parental K562 cells and *Psp*Cas13b-treated cells (parental and TKI-resistant). *Psp*Cas13b-mediated remodelling in parental and TKI-resistant cells resulted in the upregulation of genes involved in haemoglobin and heme biosynthesis, along with alterations in mitochondrial and glycolysis-related genes (**Figure 5**). This indicated that BCR::ABL1 silencing with *Psp*Cas13b mediates metabolic reprogramming, erythroid differentiation, cell cycle arrest, and apoptosis in parental and TKI-resistant K562 cells. The alterations of these gene networks by *Psp*Cas13b are consistent with previous reports using TKIs to target BCR::ABL1 in TKI-sensitive cells^57,58^, demonstrating potent suppression of oncogenic networks mediated by wildtype and T315I BCR::ABL1 variants. Thus, *Psp*Cas13b inhibits the primary oncogenic driver, overcomes common resistance to first-line TKIs, and alters downstream metabolic pathways leading to leukemic cell differentiation and apoptosis.

### Versatile platform for personalised silencing of fusion transcripts

Beyond BCR::ABL1, targeting other undruggable fusion genes has been a top priority of cancer researchers for decades due to their potent oncogenic activity in various cancers, including leukemias^44,45^, Ewing’s sarcoma^59^, and synovial sarcoma^46^. However, directly targeting these oncogenic fusions with traditional drug discovery methods, like small molecules and monoclonal antibodies, has been challenging due to structural complexity^51^. For example, despite being the most common fusion driver in AML, there is no specific drug that directly targets the RUNX1-RUNX1T1 fusion protein^44^. Our principles of personalised design of *Psp*Cas13b demonstrated its potential for potent silencing of ‘*undruggable*’ fusion transcripts including RUNX1-RUNX1T1 and EWSR1-FLI (**Figure 6**). Excitingly, the breakpoint sequence of a given fusion driver can be easily obtained from patients’ tumour RNA sequencing^60^, which would enable the manufacturing of personalised *Psp*Cas13b drugs in an actionable timeframe. The tractability and flexibility of this approach make these drug development pipelines suitable for cancer discovery as well as potential clinical translation to suppress various ‘*undruggable*’ tumour drivers within actionable time frames. Perhaps the most formidable obstacle to *Psp*Cas13b therapeutics is the lack of modalities to deliver these personalised anti-cancer drugs to tumour cells *in vivo*. Among various nanotechnologies being explored, significant advances have been made with next-generation lipid nanoparticles (LNPs) that encapsulate and deliver nucleic acid payloads to various cells and tissues including bone marrow in a targeted manner, including RNA-encoding CRISPR effectors^61,62^. These approaches may well prove to be adaptable for *Psp*Cas13b-mediated silencing of oncogenic fusion drivers in cancer cells.

In conclusion, *Psp*Cas13b design flexibility, efficacy, and specificity described here provide a new conceptual framework that could transform the landscape of personalised medicine through targeting ‘*undruggable’* or drug-resistant oncogenic RNAs including fusion transcripts.

## METHODS

### Design and cloning of crRNAs for *Psp*Cas13b

The design and cloning of *Psp*Cas13b crRNAs were according to a previous publication^20^. Briefly, individual guide RNAs were cloned into the pC0043-*Psp*Cas13b crRNA backbone (addgene#103854, a gift from Feng Zhang lab, later referred to as crRNA backbone) which contains a *Psp*Cas13b crRNA direct repeat (DR) sequence and two BbsI restriction sites for the cloning of spacer sequence. A total of 20 μg crRNA backbone was digested by BbsI restriction enzymes (NEB, R3539) following the manufacturer’s instructions for 2 hours at 37C°. Backbone linearisation was checked with 1% agarose gel. The digested backbone was purified with NucleoSpin Gel and PCR Clean-up Kit (Macherey-Nagel, 740609.50), aliquoted, and stored in -20C° until use.

For crRNA cloning, forward and reverse single-stranded DNA oligonucleotides containing CACC and CAAC overhangs respectively, were ordered from Sigma or IDT (100 μM). Forward and reverse DNA oligos were annealed in buffer (5 μl NEB buffer 3.1 and 42 μL H_2_O) by 5 min incubation at 95 °C and slowly cooled down in the heating block overnight. The annealed oligos were ligated with digested *Psp*CAs13b crRNA backbone in T4 ligation buffer (3 h, RT) (Promega, M1801).

Similarly, RfxCas13d crRNA spacers were designed as single-stranded forward and reverse DNA oligos containing AAAC and AAAA overhangs, respectively, allowing for ligation into a BsmBI-digested (NEB, R0739S) plasmid encoding a *Rfx*Cas13d direct repeat (Addgene 138150, a gift from N. Sanjana’s lab).

All crRNA spacer sequences used in this study are listed in Supplementary Table 1. All crRNAs and *Psp*Cas13b clones that were generated in this study were verified by Sanger sequencing (AGRF, AUSTRALIA). The primers used for PCR and Sanger sequencing are listed in Supplementary Table 3.

### Cloning of BCR-ABL1 mutants (T315I, C464W and T315I-C464W), BCR-ABL1, ABL1 and BCR fragments

The partial sequences of BCR-ABL1, ABL1 and BCR were designed according to the full length BCR-ABL1 P190. The IDT DNA synthesis platform provided the three sequences that were subsequently cloned into MSCV-IRES-mCherry, MSCV-IRES-eGFP and MSCV-IRES-tagBFP vectors respectively in frame with 3xHA tag using EcoRI/BamHI digestion (Promega, R6011/Promega, R6021), gel purification, and ligation with T4 DNA ligase. The ligated product was transformed into chemically competent bacteria (TOP10 or Stbl3) and positive clones were screened by PCR and Sanger sequencing (AGRF, AUSTRALIA). The BCR-ABL1-3xHA-IRES-mCherry, BCR-3xHA-IRES-tagBFP and ABL1-3xHA-IRES-EGFP constructs are available through Addgene (upon publication). The sequences of these DNA constructs are available in Supplementary File 2. The primers used for PCR and Sanger sequencing are listed in Supplementary File 3.

To generate the T315I mutation, site-directed mutagenesis was performed by using the Phusion Site-Directed Mutagenesis Kit (Thermo Fisher F541) with the following primers with 5’ phosphorylation: (T315I_F: 5’CCCCGTTCTATATCATCATCGAGTTCATGACCTAC3’ and T315I_R: 5’GCTCCCGGGTGCAGACCCCAAGGAG 3’). To generate the C464W mutation, site-directed mutagenesis was performed with the following primers with 5’ phosphorylation: (C464W_F: 5’cccagaaggctggccagagaaggtc3’ and C464W_R: 5’ cgctccatgcggtagtccttct 3’).

### Cloning of EWSR1-FLI1 fusion, EWSR1-WT and FLI1-WT constructs

Inserts for cloning were generated via PCR amplification of cDNA templates derived from fusion-positive (A673, ES8) or fusion-negative (THP-1) cell lines. Primer sets were designed to generate amplicons of ∼500bp that spanned the fusion breakpoint. Forward primers were designed to contain a Mlu1 restriction site and Kozak sequence, and reverse primers to contain a stop codon and XhoI restriction site, allowing for cloning into the MSCV-IRES-GFP reported backbone and expression of the target fragment (Supplementary Table 3). Type I EWSR1(Exon 7)::FLI1(Exon6) and Type II EWSR1(Exon 7)::FLI1(Exon5) inserts were amplified from the A673 and ES8 fusion-positive Ewing’s sarcoma cell lines, respectively. EWSR1-WT and FLI1-WT from the corresponding exonic location (i.e. EWSR1 exons 6-11 and FLI1 exons 4-9) were amplified from the fusion-negative THP1 cell line. 25 PCR cycles (with annealing temperature 60°C) were performed using Q5 High-Fidelity DNA Polymerase (NEB, M0492S) according to the manufacturer’s instructions, followed by PCR-cleanup using the NucleoSpin Gel and PCR Clean-up kit (Machery-Nagel, 740609.50). Cloning into MSCV-IRES reporter backbones encoding GFP was achieved via plasmid digestion, gel purification, and T4 ligation (Promega, M1801), and correct integration of the breakpoint (or wildtype) sequence was confirmed via Sanger Sequencing (AGRF, Australia).

### Plasmid amplification and purification

The plasmid amplification and purification are described in a previous publication^16^. Briefly, TOP10 or Stbl3 bacteria were used for transformation. A total of 5–10 μL ligated plasmids were transformed into 30 μL of chemically competent bacteria by heat shock at 42 °C for 45 s, followed by 2 min on ice. The transformed bacteria were incubated in 500 μL LB broth media containing 75 μg/mL ampicillin (Sigma-Aldrich, A9393) for 1 h at 37 °C in a shaking incubator (200 rpm). The bacteria were pelleted by centrifugation at 6,000 rpm for 1 min at room temperature (RT), re-suspended in 100 μL of LB broth, and plated onto a pre-warmed 10 cm LB agar plate containing 75 μg/mL ampicillin, and incubated at 37 °C overnight. The next day, single colonies were picked and transferred into bacterial starter cultures and incubated for ∼6 h for mini-prep (Macherey-Nagel, NucleoSpin Plasmid Mini kit for plasmid DNA, 740588.50) or maxi-prep (Macherey-Nagel**, NucleoBond Xtra Maxi Plus, 740416.50**) DNA purification according to the standard manufacturer’s protocol.

### Cell culture

The HEK 293T cell line (ATCC CRL-3216) was cultured in DMEM high glucose media (Thermo Fisher, 11965092) containing 10% heat-inactivated fetal bovine serum (Thermo Fisher, 10100147), 100 mg/mL Penicillin/-Streptomycin (Thermo Fisher, 151401220), and 2 mM GlutaMAX (Thermo Fisher, A1286001). HEK 293T cells were maintained at confluency between 20 and 80% in 37 °C incubators with 10% CO_2_. Parental K562 (ATCC CCL-243) cell line was cultured in RPMI 1640 media (Thermo Fisher, 11875083) containing 10% heat-inactivated fetal bovine serum (Thermo Fisher, 10100147) and 100mg/mL Penicillin/-Streptomycin (Thermo Fisher, 151401220). TKI-resistant (T315I) K562 cell line was cultured in RPMI 1640 media containing 200nM Dasatinib (AdipoGen Life Sciences, AG-CR1-3540)^39^, 10% heat-inactivated fetal bovine serum and 100mg/mL Penicillin/-Streptomycin. K562 cells were maintained at confluency between 20 and 80% in 37 °C incubators with 5% CO_2_. Cells were routinely tested and were mycoplasma-negative.

### RNA silencing assays using plasmid transfections

All transfection experiments were performed using an optimised Lipofectamine 3000 transfection protocol (Thermo Fisher, L3000015)^22^. For RNA silencing in HEK 293T, cells were plated at approximately 30,000 cells/100 μL/96-well in tissue culture-treated flat-bottom 96-well plates (Corning) 18 h prior to transfection. For each well, a total of 100 ng DNA plasmids (22 ng of *Psp*Cas13b-NES-3xFLAG-T2A-BFP (addgene #173029) or pC0046-EF1a-*Psp*Cas13b-NES-HIV (addgene #103862), 22 ng crRNA plasmid, and 56 ng of the target gene plasmid) were mixed with 0.2 μL P3000 reagent in Opti-MEM Serum-free Medium (Thermo Fisher, 31985070) to a total of 5 μL (mix1). Separately, 4.7 μL of Opti-MEM was mixed with 0.3 μL Lipofectamine 3000 (mix2). Mix1 and mix2 were added together and incubated for 20 min at room temperature, then 10 μL of transfection mixture was added to each well. Supplementary Table 4 summarises the transfection conditions used in 96, 24, and 12-well plates. After transfection, cells were incubated at 37C°, 10% CO_2_, and the transfection efficacy was monitored 24-72 hours post-transfection by fluorescent microscopy.

### Cell Nucleofection

All RNA silencing experiments in K562 cells were performed using optimised SF cell line 4D-Nucleofector X Kit S protocol (Lonza, V4XC-2032). For each reaction, 1×10^6^ cells were resuspended in 20 μL SF Cell Line 4D-Nucleofector™ X Solution containing 3.5 μg crRNA plasmid. The K562 cells were nucleofected using program FF-120, 4D-Nulceofector X unit (Lonza, AAF-1003X) on Day 1. Cells were placed in the incubator *overnight for recovery and crRNA expression. On day 2, K562 cells were* nucleofected again using 20 μL of SF Cell Line 4D-Nucleofector X Solution containing 3.5 μg Cas13-BFP plasmids. Alternatively, 1×10^6^ cells were resuspended in 20 μL SF Cell Line 4D-Nucleofector™ X Solution containing 2 μg crRNA plasmid and 4 μg Cas13-mFlame mRNA or Cas13 (untagged) mRNA (Messenger Bio). Cells were incubated at 37 °C and 5% CO_2_ for 48 to 96 hours, and the silencing efficiency was monitored by RT-PCR, western blot, or FACS analysis.

### Fluorescent microscopy analysis

For RNA silencing experiments, the fluorescence intensity was monitored using the EVOS M5000 FL Cell Imaging System (Thermo Fisher). Images were taken 48 h post-transfection, and the fluorescence intensity of each image was quantified using a lab-written macro in ImageJ software. Briefly, all images obtained from a single experiment are simultaneously processed using a batch mode macro. First, images were converted to 8-bit, threshold adjusted, converted to black and white using the Convert to Mask function, and fluorescence intensity per pixel measured using the Analyze Particles function. Each single mean fluorescence intensity was obtained from four different field of views for each crRNA, and subsequently normalised to the non-targeting (NT) control crRNA. A two-fold or higher reduction in fluorescence intensity is considered biologically relevant.

### Western Blot

Cells were washed three times with ice-cold PBS ± and lysed on ice in RIPA lysis buffer [50 mM Tris (Sigma-Aldrich, T1530), pH 8.0, 150 mM NaCl, 1% NP-40 (Sigma-Aldrich, I18896), 0.1% SDS, 0.5% sodium deoxycholate (Sigma-Aldrich, D6750)] containing protease inhibitor cocktail (Roche, 04693159001) and phosphatase inhibitor cocktail (Roche, 4906845001). Samples were incubated for 30min at 4 °C with rotation (25 rpm), and centrifuged at 16,000 g for 10 min, 4 °C. Supernatant was transferred to a new tube. Protein concentrations were quantified using the Pierce BCA Protein Assay Kit (Thermo Fisher, 23225) according to the manufacturer’s instructions. A total of 10 μg of protein diluted in 1x Bolt LDS sample buffer (Thermo Fisher, B007) and 1x Bolt sample reducing agent (Thermo Fisher, B009) were denatured at 95 °C for 5 min. Samples were resolved by Bolt Bis-Tris Plus 4–12% gels (Thermo Fisher, NW04120BOX) in 1x MES SDS running buffer (Thermo Fisher, B0002) and transferred to 0.45 μM PVDF membranes (Thermo Fisher, 88518) by a Trans-Blot Semi-Dry electrophoretic transfer cell (Bio-Rad) at 20 Volt for 30 min. Alternatively, samples were resolved by 4-15% Criterion TGX Precast Midi Protein gels (Bio-Rad, 5671084) in 1x Tris/glycine/SDS running buffer (Bio-Rad, 1610732) and transferred to 0.20 μM nitrocellulose membranes (Bio-Rad, 1704159) by a Trans-Blot Turbo Transfer System (Bio-Rad) with a HIGH MW protocol. Membranes were incubated in blocking buffer 5% (w/v) BSA (Sigma-Aldrich, A3059) in TBST with 0.15% Tween 20 (Sigma-Aldrich, P1379) for 1h at RT and probed overnight with primary antibodies at 4 °C. Blots were washed three times in TBST with 0.15% Tween20, followed by incubation with fluorophore-conjugated or HRP-conjugated secondary antibodies for 1h at RT. Membranes were washed in TBST (0.15% Tween20) three times and fluorescence or chemiluminescence was detected using the Odyssey CLx Imager 9140 (Li-cor), iBright CL1500 Imaging System (Thermo Fisher), or ChemiDoc Imaging System (Bio-Rad). The antibodies used for western blots are listed in Supplementary Table 5. Resources Table.

### Cell flow cytometry

*For intracellular staining, ∼*1×10^6^ *cells were fixed with Fixation Buffer (BioLegend, 420801) and permeabilised with True-Phos Perm Buffer (BioLegend, 425401) according to the manufacturer’s instructions.* Samples were then incubated in 50 µL PBS^-/-^/2% (v/v) FBS containing primary antibody for 1 hour at RT, followed by washing twice with 200µl PBS^-/-^ 2% (v/v) FBS. Next, samples were incubated with 50 µL PBS^-/-^/2% (v/v) FBS containing an Alexa fluor 647 goat anti-rabbit IgG (H+L) secondary antibody (CAT# A21245) for 30 minutes at RT, followed by washing twice with 200µl PBS^-/-^ /2% (v/v) FBS. Finally, cells were re-suspended in 200 μL of 1x PBS containing 2% FBS for flow cytometry analysis. All samples were analysed by FACSymphony Cell Analyzer A5 or A3 (BD Biosciences).

For cell sorting, cells were resuspended in PBS^-/-^/2% (v/v) FBS containing 1 ng/mL PI in 5 ml FACS tubes, sorted using a BD FACSAria Fusion 5 or Fusion 3 instrument and re-collected in fresh media for RT-PCR, western blots or proliferation assays.

All flow cytometry profiles were analysed using FlowJo V10 software (Tree Star Inc). The antibodies used for cell flow cytometry are listed in Supplementary Table 5.

### Resazurin-based proliferation assay

Resazurin reagent was used to quantify K562 cell proliferation. Resazurin reagent contains 75 mg of Resazurin (Sigma, R7017), 12.5 mg of methylene blue (Sigma, MB-1), 164.5 mg of potassium hexacyanoferrate (III) (Sigma, P8131), and 211 mg potassium hexacyanoferrate (II) trihydrate (Sigma, P9387), resuspended in 500 mL PBS. On Day 0, 10,000 K562 cells were seeded in 200 μL fresh media per well in a 96-well plate for Rezazurin proliferation assay with technical triplicate per condition. 40 μL of resazurin reagent was added to each well and incubated at 37 °C and 5% CO_2_ for 4 hours. The absorbance was measured at time points 0, 24, 48, 72, and 96 hours using Cytation3 Plate Reader (BioTek, wavelengths 550 nm and 590 nm). 1 µM of *Imatinib* was used as a positive control to inhibit BCR::ABL1 and K562 proliferation, while 0.01% of DMSO was used as vehicle treatment.

### RNA extraction, cDNA synthesis, RT-PCR, and PCR

Total RNA was isolated from ∼ 0.2-1×10^6^ cells using the NucleoSpin RNA Plus (Macherey-Nagel, 740984.50) or *Quick-*RNA Miniprep Kit (*Zymo* Research, R1055) following the manufacturer’s instructions. Alternatively, total RNA was isolated by standard Trizol-chloroform extraction according to manufacturer’s instruction (Thermo Fisher, 15596026), followed by Dnase treatment with RQ1 RNase-Free DNase according to manufacturer’s instruction (Promega, M6101). RNA integrity was analysed by Agilent 2200 Tapestation (Agilent, G2964AA) with RNA ScreenTape (Agilent, 5067-5576) according to the manufacturer’s instructions.

1μg total RNA was converted to cDNA using the high-capacity cDNA reverse transcription kit (*Thermo Fisher,* 4368814) following the manufacturer’s instructions.

Quantitative RT-PCR reaction was performed in a technical duplicate using the StepOne Real-Time PCR system (*Thermo Fisher*) and PowerUp™ SYBR™ Green Master Mix (*Thermo Fisher*, A25742). The reaction contains 0.2 μL of cDNA, 0.6μM of forward primer, and 0.6 μM of the reverse primer.

The PCR reaction was performed using a Q5 High-Fidelity DNA kit (NEB, E0555S). The reaction contains 1 μL cDNA, 0.5 μM forward primer, and 0.5 μM reverse primer*. The PCR product was purified with* NucleoSpin Gel and PCR Clean-up kits (Macherey-Nagel, 740609.50). PCR products were evaluated using 1% agarose gel electrophoresis. Primers for RT-PCR and PCR are detailed in Supplementary Table 3.

### Whole Transcriptomic RNA sequencing

Total RNA was isolated using the NucleoSpin RNA Plus (Macherey-Nagel, 740984.50) following the manufacturer’s instructions. RNA was measured by Qubit assays to determine dsDNA contamination. RNA was processed by the Molecular Genomics Facility (Peter MacCallum Cancer Centre) or Australian Genome Research Facility (AGRF) using Illumina Ribo-Zero Plus rRNA Depletion Kit and pair-end sequenced on an Illumina NextSeq 500 or NovaSeq 6000 System. Libraries (triplicate RNA samples) were created from the samples and combined for sequencing with 150bp reads, for a total of ∼4 to 7 million reads per HEK 293T cell sample and for a total of ∼50 million reads per k562 cell sample. The raw data files were converted to FASTQsanger and mapped to the modified Human reference transcriptome (GRCh37 ver.19) with BCR::ABL1 RNA sequences added using the Galaxy platform^63^. Subsequently, fusion transcripts were detected using Arriba^64^. Indexing of BAM files was performed in samtools for alignment visualisation using IGV.

Genes without more than 1 CPM (counts per million) for HEK 293T cell samples or 2 CPM for K562 cell samples in at least 3 samples were considered to be insignificant and filtered out. Read counts were normalised for effective library size and differential expression analysis between targeting and non-targeting samples was performed using the voom-limma (v3.42.2) pipeline^65^. Genes with log_2_ fold change (FC) >1 were considered upregulated while genes with log_2_ FC <-1 were considered downregulated. Adjusted *P*-value (with 0.05 false discovery rate) < 0.05 was considered statistically significant. Transcripts of interest or transcripts that were significantly up or down-regulated between targeting and NT crRNAs are highlighted in the volcano plots.

For the breakpoint read analysis, the alignment was re-performed using STAR (v2.7.10b) against GRCh38, including an additional genetic element corresponding to either the HEK293T or K562 BCR::ABL1 fusion gene. Subsequent analysis was performed in R (v4.2.2). Read counts were normalised across samples utilising counts per million (CPM). SAM file alignments were filtered for those mapping exclusively to the fusion gene, discarding any multi-mapping read alignments whilst allowing a sequence mismatch of 1. For the HEK 293T cell data where the exogenously expressed BCR::ABL1 contained no introns, breakpoint reads, defined as matched single reads with 5’ and 3’ ends mapping respectively to BCR and ABL1, were identified based on read start and end positions. Non-canonical breakpoint split-reads – defined as breakpoint reads containing at least one gap ≥ 20 nucleotides – were identified via CIGAR strings. This was similarly performed to identify non-canonical split-reads across the whole fusion gene (Supp. Figure 2d), except, in this instance, all alignments to BCR::ABL1 were used, allowing for multi-mappers to endogenous BCR and ABL1. For the K562 data where BCR::ABL1 is endogenously expressed, canonical breakpoint reads were identified via the correct corresponding splicing over the fusion region based on the reference annotation. Breakpoint reads without this correct splicing profile were assigned as non-canonical breakpoint split-reads.

All transcriptomic data are provided with the paper in a source data file.

### Samples processing for mass spectrometry analysis

48h post-transfection, cells were washed three times with ice-cold PBS -/- and lysed on ice in RIPA lysis buffer [50 mM Tris (Sigma-Aldrich, T1530), pH 8.0, 150 mM NaCl, 1% NP-40 (Sigma-Aldrich, I18896), 0.1% SDS, 0.5% sodium deoxycholate (Sigma-Aldrich, D6750)] containing protease inhibitor cocktail (Roche, 04693159001) and phosphatase inhibitor cocktail (Roche, 4906845001). 10 mM of TCEP (Tris(2-carboxyethyl)phosphine) (Sigma-Aldrich, C4706) were added to 50 µg of protein lysates and incubated at 65 °C for 20 min for disulfide bond reduction. Iodoacetamide (Sigma-Aldrich, A3221) was added to a final concentration of 55 mM to alkylate proteins, samples were incubated at 37°C for 45 min in the dark. Samples were acidified by the addition of 2.5% final concentration of phosphoric acid. Then, S-Trap binding buffer [90% MeOH, 100 mM Triethylammonium bicarbonate (TEAB, Sigma-Aldrich, T7408), pH 7.1] was added to the acidified lysate, and samples were passed through Suspension trapping (S-trap) column (PROTIFI, K02MICRO10). The bound samples were then washed three times with the S-Trap binding buffer. The bound proteins were then digested overnight at 37°C by trypsin (Thermo Fisher, 90058) in 50 mM TEAB at a 1:10 ratio. The peptides were eluted with 40 µL of 50 mM TEAB, followed by 40 µL 0.2% aqueous formic acid, and finally with 35 µL of 50% acetonitrile containing 0.2% formic acid. Samples were dried in a SpeedVac concentrator (Eppendorf Concentrator Plus), resuspended in 20 µL 2% acetonitrile, 0.1% trifluoroacetic acid, sonicated for 10 min at room temperature, centrifuged at 16,000 × *g* for 10 min, and transferred into HPLC vials for analysis.

### Mass spectrometry proteomics analysis

Samples were analysed by LC-MS/MS using Orbitrap Exploris 480 (Thermo Scientific) fitted with nanoflow reversed-phase-HPLC (Ultimate 3000 RSLC, Dionex). The nano-LC system was equipped with an Acclaim Pepmap nano-trap column (Dionex – C18, 100 Å, 75 μm × 2 cm) and an Acclaim Pepmap RSLC analytical column (Dionex – C18, 100 Å, 75 μm × 50 cm). Typically for each LC-MS/MS run, 1 μL of the peptide mix was loaded onto the enrichment (trap) column at an isocratic flow of 5 μL/min of 2% Acetonitrile containing 0.05% trifluoroacetic acid for 6 min before the analytical column is switched in-line. The buffers used for the LC were 0.1% v/v formic acid in water (solvent A) and 100% Acetonitrile/0.1% formic acid v/v (Solvent B). The gradient used was 3% B to 23% B for 89 min, 23% B to 40% B in 10 min, 40% B to 80% B in 5 min and maintained at 80% B for the final 5 min before equilibration for 10 min at 2% B prior to the next analysis. All spectra were acquired in positive mode with full scan MS spectra scanning from m/z 300-1600 at 120,000 resolution with AGC target of 3×10^6^ with a maximum accumulation time of 25 ms. The peptide ions with charge state ≥2-6 were isolated with an isolation window of 1.2 m/z and fragmented with a normalised collision energy of 30 at 15,000 resolution.

All generated files were analysed with MaxQuant (version2.2.0.0) and its implemented Andromeda search engine to obtain protein identifications as well as their label-free quantitation (LFQ) intensities. Database searching was performed with the following parameters: cysteine carbamidomethyl as a fixed modification; oxidation and acetyl as variable modifications with up to 2 missed cleavages permitted; main mass tolerance of 4.5 ppm; 1% protein false discovery rate (FDR) for protein and peptide identification; and minimum two peptides for pair-wise comparison in each protein for label-free quantitation. The raw data files were searched against the modified Human reference proteomes (UP000005640).

The MaxQuant result output was further processed with Perseus (Version 2.0.7.0), a module from the MaxQuant suite. After removing reversed and known contaminant proteins, the LFQ values were log_2_ transformed and the reproducibility across the biological replicates was evaluated by a Pearson’s correlation analysis. The replicates were grouped accordingly, and all proteins were removed that had less than 4 “valid values” in each group. The missing values were replaced by imputation, and differential expression analysis between targeting and non-targeting samples was performed using a two-sample *t*-test. Proteins with log_2_ fold change (FC)>1 were considered upregulated while proteins with log_2_FC<-1 were considered downregulated. *P*-value < 0.05 was considered as statistically significant. Proteins of interest or proteins that were significantly up or down-regulated between targeting and NT crRNAs are highlighted in the volcano plots. All proteomics data are provided with the paper in a source data file.

### Data analysis

Data analyses and visualisations (graphs) were performed in GraphPad Prism software version 9, unless stated otherwise. In some figures, identical fluorescence data points from an NT control are displayed in multiple panels to facilitate comparison with the targeting condition. The figures displaying identical NT data points are listed in the Source Data for reference. Specific statistical tests and numbers of independent biological replicates are mentioned in respective figure legends. The silencing efficiency of various crRNAs was analysed using an unpaired two-tailed Student’s *t*-test or one-way analysis of variance (ANOVA) followed by Dunnett’s multiple comparison test where we compared every mean to a control mean as indicated in the figures (95% confidence interval). The *P* values are indicated in the figures. *P* < 0.05 was considered statistically significant. Pearson’s correlation coefficient was used to analyse the correlation between the log_2_ fold changes of genes detected by RNA sequencing and mass spectrometry. Protein-protein interaction network analyses were performed by STRING^66^. Gene ontology analyses were performed by DAVID and ShinyGO^67,68^. The Chord ontology plot and the Sakey ontology plot were generated by SRplot^69^.

## Data availability

All data are available in the main text and supplementary materials. Source Data will be available on Figshare once published. All key plasmids constructed in this study, their sequences, and maps will be deposited to Addgene upon publication. Reference human proteome (UP000005640) and reference human transcriptome (GRCh37 ver.19) are used in this study. The mass spectrometry proteomics data will be deposited to the ProteomeXchange Consortium via the PRIDE partner repository with the dataset identifier available upon publication. The RNA sequencing transcriptomic data will be deposited to the NCBI’s Gene Expression Omnibus (GEO) with the dataset identifier available upon publication.

## Acknowledgments

The authors thank all lab members for facilitating experiments and discussions.

## Funding

This work was supported by a Cancer Council Victoria Ventures grant (829606 to MF, PGE, and JAT); mAP mRNA Victoria grant (RCH0153742 to MF); the CASS foundation, and the Australian and New Zealand Sarcoma Association (ANZSA).

## Notes

### Competing Interest Statement

The authors have declared no competing interest.

